# Serum-free media development and validation for cultivation of C2C12 immortalised murine myosatellite cell line for cultivated meat

**DOI:** 10.64898/2026.07.06.736713

**Authors:** William Gordon-Petrovskii, Maria Laura Vieri, Benjamin AS Dagès, Michael Sulu, Isa Senica, Gary J Lye, Mariana Petronela Hanga

## Abstract

The development of cost-effective, serum-free media is critical for scalable cultivated meat production. This study used high-throughput screening through a Design of Experiments (DoE) approach to develop an animal-free, serum-free medium (MMM1) specifically for the C2C12 murine myoblasts model cell line with applicability in cultivated meat research including for pet food. Low cost, food-grade inputs such as methylcellulose and spirulina extract resulted in significant cell growth improvements. The optimised MMM1 formulation containing low cost, food-grade inputs, achieved cumulative population doublings comparable to 10% (v/v) fetal bovine serum over four consecutive passages. Furthermore, MMM1 supported scalable cell expansion on commercially available dextran-based microcarriers (Cytodex-3) in both static and agitated conditions in spinner flasks, matching growth rates of serum-based controls. Finally, transitioning to a food-grade DMEM/F12 basal medium maintained cell proliferation equivalent to the pharmaceutical-grade DMEM/F12, but at a significantly lower cost, thus offering a viable strategy to substantially reduce biomanufacturing costs which is a critical challenge in cultivated meat production.

**Highlights:**

- A serum-free medium formulation for C2C12 murine myoblasts (MMM1)
- C2C12 growth in MMM1 comparable to serum-based medium
- C2C12 growth in MMM1 in microcarrier culture as effective as serum-based medium
- MMM1 can be translated to be animal-free and fully food-grade

**Graphical abstract:** 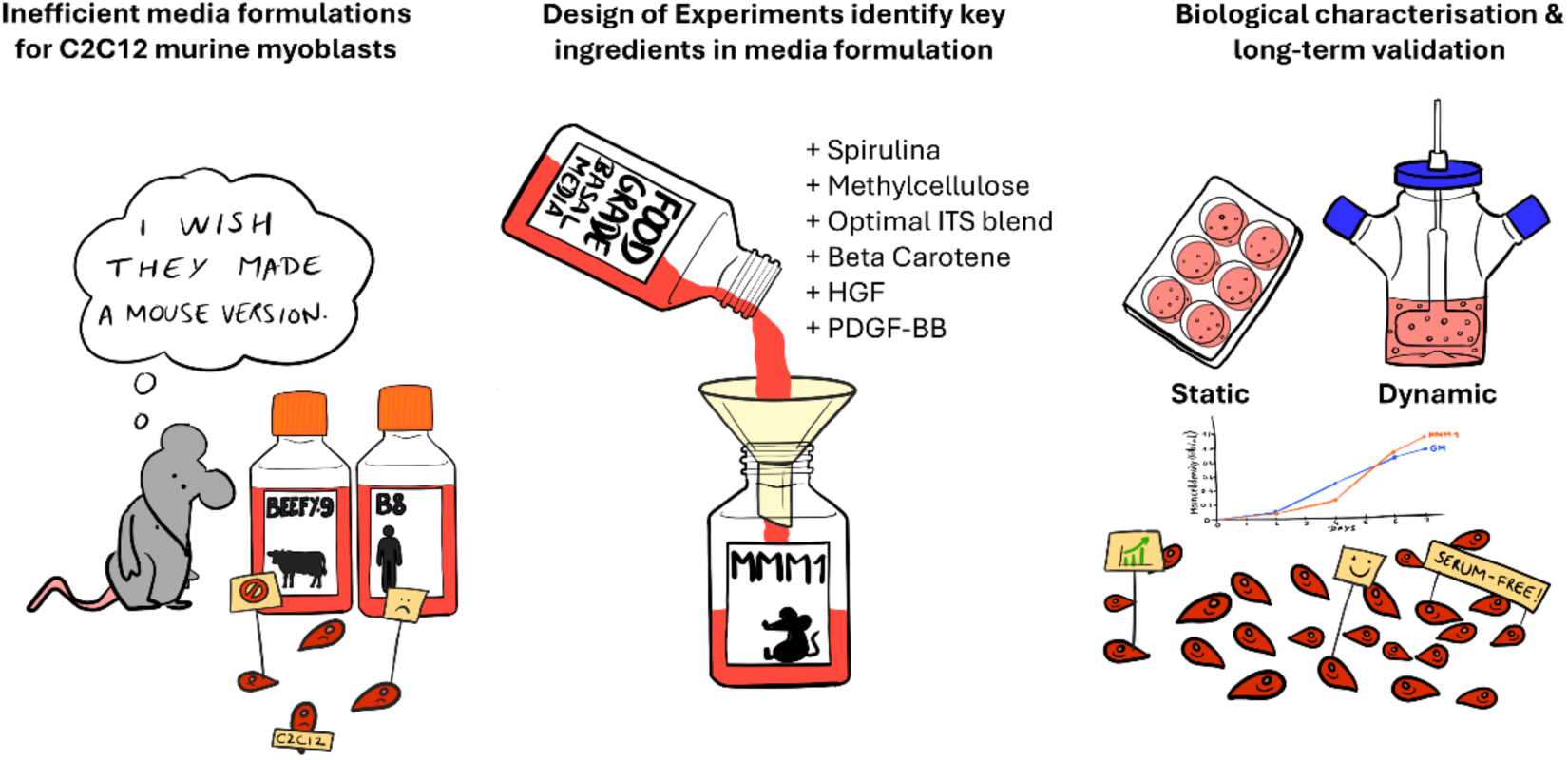

## 1. Introduction

Cultivated meat is a novel food technology for meat production through the *in vitro* cultivation of animal cells (Post, 2014). This novel food has the potential to help curb the impacts of increased projected global meat consumption (Font-i-Furnols, 2023). Cultivated meat production offers distinct advantages over traditional livestock farming, specifically regarding resource efficiency and public health. Research indicates that cultivated beef, in particular, requires a fraction of the land needed for conventional agriculture (Sinke et al., 2023). Furthermore, the process largely eliminates the necessity for antibiotics (McNamara & Bomkamp, 2022) unlike industrialised animal farming. Beyond these technical metrics, cultivated meat also resolves the profound animal welfare concerns associated with industrial rearing and slaughter (Huber, 2024; Stefanutti, 2024).

Traditional animal cell culture practices often rely on foetal bovine serum (FBS) as an effective cell culture supplement for a variety of cell types (D. Y. Lee et al., 2022). However, FBS is largely unsuitable for the production of cultivated meat. First and foremost, because it is expensive, costing ∼$500-1,000 per litre. Furthermore, being animal derived, it limits the potential positive impacts of cultivated meat regarding reliance on animals. Therefore, there is a need for an alternative serum-free medium, with minimal to no reliance on animal components, and based on sustainable components.

To achieve industrial viability of cultivated meat, removal of serum is only the first step. Food-grade inputs will also play a pivotal role in achieving compliance with food regulations, lowering manufacturing costs to achieve cost parity with traditional meat, and ensuring supply chains are available for high production volumes. Switching all media components from pharmaceutical-grade to food-grade could reduce costs up to 95% (Specht, 2020). Significant progress has been made to date in the development of serum-free media for cultivated meat, resulting in formulations such as B8 (Kuo et al., 2020), Beefy-9 (Stout, Mirliani, et al., 2022) and others (Cosenza et al., 2022; Kolkmann et al., 2020; Lawson & Purslow, 2000; Messmer et al., 2022; Pasitka et al., 2022; Skrivergaard et al., 2023). However, serum-free media formulations are cell line-specific and as such, when used with other cells than the ones that it was optimised for, the cell performance can be significantly different (Kolkmann et al., 2020; Tan et al., 2015) For example, Kolkmann et al (2020) tested 7 commercially available serum-free media formulations with primary bovine myoblasts with only 3 formulations resulting in cell growth, but only for 6 days.

The most popular choice of starting cells for production of cultivated meat was found to be myosatellite cells as detailed in a 2023 industry survey carried out by The Good Food Institute (Ravikumar et al., 2024). Myosatellite cells are unipotent stem cells that physiologically play a key role in muscle repair and regeneration through their ability to differentiate into myotubes (Yin et al., 2013). Due to the limited availability of commercially available myosatellites from relevant species for cultivated meat production (e.g. bovine, porcine, avian etc), many research groups have chosen to use the commercially available C2C12 murine immortalised satellite cell line (Yaffe & Saxel, 1977) as a model cell line for cultivated meat research and promising cell line for cultivated pet food research. The C2C12 cell line possesses many of the industry’s most desired traits for cultivated meat cell lines (Ravikumar et al., 2024) including fast proliferation rate with a population doubling time of ∼19.4 hours (Humphrey et al., 2012), spontaneous immortalisation (Yaffe & Saxel, 1977) and strong targeted differentiation potential (Bajaj et al., 2011). Additionally, cultivated meat also includes pet food (Stefanutti, 2024) with two regulatory approvals to date, one in the UK (H. Jones, 2024) and one in Singapore (Mridul, 2025). With these traits, C2C12 can be considered a robust model cell line for cultivated meat that can be used to study both upstream and downstream processing.

This study uses high-throughput screening through a Design of Experiments (DoE) approach to develop an animal-free, serum-free medium (MMM1) specifically for the C2C12 murine myoblasts model cell line with applicability in cultivated meat research including for pet food. While serum-free media formulations for myosatellite cells have been previously published (Stout, Mirliani, et al., 2022, Cosenza et al., 2022, Kolkmann et al, 2020, Pasitka et al., 2022; Skrivergaard et al., 2023), those formulations rely on high concentrations of growth factors and proteins such as albumin or fetuin. In this study, we took the approach of exploring lower-cost, food grade components such as spirulina, methylcellulose and beta carotene with the aim to lower costs and lower the reliance on expensive proteins.

## 2. Materials and methods

### 2.1 Materials and reagents

Growth factors and other supplements were prepared according to the manufacturers’ recommendation unless otherwise stated. The following materials and reagents were used throughout this study: Dulbecco’s Modified Essential Medium/F12 (DMEM/F12) basal medium (Gibco 11320033), Food-Grade DMEM/F12 (Multus, DMFG01), fetal bovine serum (FBS) (heat-inactivated, South American origin, Thermofisher, 10270106), Tryple Select (Gibco, 12604021), dimethyl sulfoxide (DMSO) (Sigma, D4540), horse serum (Gibco, 16050130), Presto Blue reagent (Invitrogen, A13262), paraformaldehyde (J19943.K2, Thermofisher Scientific), Hoechst 33342 (Thermofisher, H3570), L-ascorbic acid 2-phosphate (Merck, 49752), insulin-transferrin-selenium (ITS -G) (100X) (Gibco, 41400045), insulin-transferrin-selenium-ethanolamine (ITS -X) (100X) (Gibco, 51500056), human recombinant insulin (Merck, I0908), human recombinant transferrin (Merck, T3705), human recombinant FGF2 (Peprotech, 100-18B), sodium selenite (Sigma-Aldrich S5261), human recombinant neuregulin-1 (NRG1) (MedChemExpress, HY-P7365), human recombinant transforming growth factor – beta (TGFβ-1) (Peprotech, 100-21), recombinant human albumin (Merck, A9731), recombinant human hepatocyte growth factor (HGF) (BioLegend, 596402), recombinant human interleukin-6 (IL-6) (BioLegend, 570804), bovine serum albumin (BSA) (Sigma, A2153), recombinant human platelet-derived growth factor BB (PDGF-BB) (BioLegend, 577302), recombinant human insulin-like growth factor 1 (IGF-1) (Cayman Chemical, CAY33981), linoleic acid (CAY90150), hydrocortisone (Thermo Scientific, 10062031), L-carnosine (Merck, C9625), curcumin (Abcam, ab120618), beta carotene (Stratech, S1767-SEL), methylcellulose (Special Ingredients, 5060341112959), guar gum (Special Ingredients, 5060341113185), xanthan gum (Special Ingredients 5060341110511), DL-alanine (Merck, A7502), dextran (Clinical grade mW 60,00-90000; MP Biomedicals 180140), City-mix: Shanghai Mix (3D Bio-Tissues, CM-Sh-15), Singapore Mix (3D Bio-Tissues, CM-Sg-15), London Mix (3D Bio-Tissues, CM-Ln-15), Miami Mix (3D Bio-Tissues, CM-Mi-15), Tokyo mix (3D Bio-Tissues, CM-Ty-15).

### 2.2. Spirulina extract preparation

Spirulina extracts were prepared using a protocol developed in Gordon-Petrovskii et al. (2026), by suspending the commercial product to create a 5% (w/v) solution in ultra-pure water. The resulting solution was then subjected to ultrasonication on ice for 24 cycles, consisting of 10 seconds of active sonication followed by 10 seconds of rest. Following sonication, the mixture was centrifuged at 8000 xg to pellet the insoluble cellular debris. The resulting supernatant was collected and sterilised by filtration through a 0.22 µm filter to yield the final working extract.

To evaluate the impact of raw material variability on cell proliferation, three distinct commercial lots of spirulina were prepared and tested. These included two independent batches sourced from Naturya (Lot W04057 and Lot W06466) and one batch from Bulk Powders (Lot 5550000052527). For this comparative batch-to-batch analysis, each sterile extract was supplemented into the formulated medium at a final concentration of 0.02% (v/v), cultured for 4 days before counting.

### 2.3 Routine monolayer cell culture

C2C12 cells were purchased from Merck (91031101) and were propagated on standard, uncoated tissue-culture treated vessels in a humidified incubator at 37°C with 5% CO₂. The Growth Medium (GM) consisted of DMEM/F12 enriched with 5 ng/mL human recombinant FGF2 and 20% (v/v) heat-inactivated FBS. For serum-free media formulation experiments, a control medium consisting of DMEM/F12 supplemented with 10% (v/v) FBS was used. Cell subculturing was performed using 1x TrypLE Select when cells reached approximately 70% confluence as observed by phase contrast microscopy. For long-term storage, cells were cryopreserved in FBS supplemented with 10% (v/v) DMSO. For differentiation to the myogenic lineage, cells were seeded at 30,000 cells/cm^2^ in growth medium for 24h to allow cell attachment, after which the medium was changed to differentiation medium (DM) consisting of DMEM/F12 with 2% (v/v) horse serum. Myogenic differentiation was carried out for 7-9 days with medium changes every 2 days.

Cell counts and viability were assessed using the Chemometec NC-3000 automated cell counter with the Via1 cassette (Chemometec, 941-0012) and protocol ‘Cell Counting and Viability’. Cell morphology was monitored using a phase contrast microscope (Evos XL, Thermofisher).

### 2.4 Multiple passage assay

C2C12 cell line is an immortalised cell line and as such, we hypothesised that there is little variation in cell growth kinetics from passage to passage. To verify this hypothesis, C2C12 cells were seeded at 300 cells/cm^2^ and subcultured after a fixed duration of 7 days for 4 consecutive passages between passages 25 and 28.

A multiple passage approach was also taken to validate the developed serum-free media formulations. For these experiments, a confluency-based subculturing methodology was adopted to account for different growth rates due to the different media formulations tested. C2C12 cells were seeded at 5,000 cells/cm^2^ with media exchanged every 2 days. For long-term,8 passage validation a lower seeding density of 2500 cells/cm^2^ to ensure a consistent passaging schedule for both mediums could be performed. Upon reaching 70% confluency in any of the conditions tested as observed by phase contrast microscopy, the cells in all conditions were harvested, and counted. While the C2C12 line is highly stable and spontaneously immortalised rendering minor passage variances typically negligible compared to primary cells, the slight passage offset between the baseline characterisation (P25–28) and this validation phase (P30–32) is acknowledged as an operational limitation. Late-passage phenotypic drift remains an inherent risk when extrapolating these kinetics to extended seed-train simulations. Population doubling time (h), population doubling level (PDL), and fold change were calculated using equation (1) (R. H. Lee et al., 2004), equations (2&3) (Cristofalo et al., 1998) and equation (4) (Hanga et al., 2020) respectively.

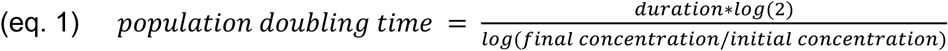

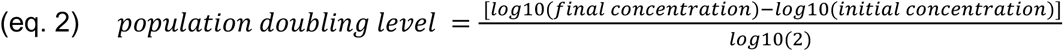

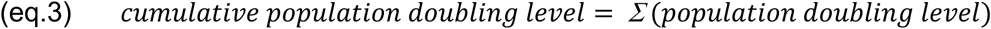

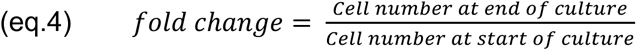

### 2.5 Microcarrier cell culture

The serum-free media formulations were validated in both monolayer (multiple passage methodology) and scalable microcarrier cultures. Two types of commercially available microcarriers were selected and used in this study. Cytodex-3 (Cytiva) and Cultispher-S (Percell Biolytica) were selected. Cytodex-3 is a microporous, dextran-based microcarrier that is positively charged and collagen coated (Rozwadowska et al., 2016), while Cultispher-S is macro-porous, gelatine-based, with no surface charge or coating (Yang et al., 2022). These two microcarriers were chosen as they provide two very different surface chemistries.

The microcarriers were tested in both static and dynamic conditions. Static cultures were carried out in ultralow attachment well plates (Corning) to ensure cell attachment only to microcarriers rather than the culture plate, while dynamic cultures were carried out in 100 mL glass spinner flasks with a paddle blade impeller (Bellco). For static culture, a concentrated stock of microcarriers was produced by weighing the corresponding dry mass, rehydrating it for 1 hour in PBS (without Ca and Mg) and then sterilising it by autoclaving. The concentrated stock was either stored at 4°C for later use or used immediately. For dynamic cultures, microcarriers were weighed, added directly to a spinner flask containing PBS and then autoclaved. To avoid cell attachment to the glass walls of the spinner flask, Sigmacote (Sigma-Aldrich, SL2-25ML) was used to siliconise the glass vessel. The silicone coating was refreshed after three uses.

Cytodex 3 microcarriers were used at a density of 5 cm^2^/mL, while Cultispher-S at 1 mg/mL. Spinner flasks were operated at a constant stirring speed of 30 rpm. 80% medium exchanges were performed at days 3, 5 and 6. Cell-microcarrier samples were taken regularly from spinner flasks for direct counting performed on the Nucleocounter NC3000 (Chemometec) using the ‘Count of Aggregated Cells A100 and B Assay’ protocol. Briefly, the cells on microcarriers were first lysed using an equal volume of Reagent A100 and then stabilised using an equal volume of Reagent B. The resulting suspension was then loaded onto Via-1 cassettes and counted.

#### 2.5.1 Metabolite & Nutrient Analysis

Spent media samples were collected and stored at -80°C for later analysis. When ready for analysis, samples are thawed and centrifuged for 10 minutes at 5000 xg. Samples are then analysed for concentrations of glucose, ammonia and lactate concentrations in the Optocell CuBiAn VC biochemistry analyzer (4BioCell; Bielefeld, Germany).

### 2.6 High-Throughput Screening

A high-throughput (HT) screening platform was established to assess the effect of different media formulations and supplementation with different components. This workflow used 96-well microplates imaged on the automated high-content spinning-disk confocal imaging system Opera Phenix (Revvity) and analysed using two different methods: nuclei quantification (2.6.1) and cell metabolic activity (2.6.2). This parallel approach allows for the simultaneous evaluation of multiple factors and concentration ranges defined by various Design of Experiments (DoE) matrices generated.

Working cell banks of C2C12 cells (passage 19) were thawed and cultured for 6 days into GM prior to any media development experiments. This approach was taken to ensure all experiments used cells at the same stage of the growth cycle and they have recovered from thawing. For media development experiments, the cells were seeded into 96-well plates at a density of 2,500 cells/cm². Media exchanges were performed on days 1 and 3, with cell growth assessed on day 4 by the two selected methods.

#### 2.6.1 Nuclei quantification

The cells were fixed using 4% (v/v) paraformaldehyde for 10 minutes at room temperature. Nuclei were stained using a Hoechst 33342 Solution (4 µM in PBS) for 10 minutes, then imaged on the Opera Phenix High throughput cell imaging system using the 5x air objective in non-confocal mode in the Hoechst 333342 channel at 100 ms excitation. Nuclei counts were obtained from the Columbus software using the “count nuclei” protocol.

#### 2.6.2 Presto Blue assay

In parallel to the Hoechst assay, Presto Blue assay was also chosen as it can be easily translated to automated platforms. Presto Blue was conducted following the manufacturer’s instructions. Briefly, a 10% (v/v) solution of Presto Blue reagent prepared in growth media was added to the cells and incubated for 1 hour at 37°C in a 5% CO_2_ humidified incubator. Samples and controls were transferred in triplicates to black-walled, clear-bottomed tissue culture plates and fluorescence was measured at 570 nm and 600 nm on a BMG Labtech Clariostar® microplate reader. The blank was growth media alone, while the negative control was fresh 10% (v/v) Presto Blue solution (unreduced) and the positive control was a 100% reduced form of Presto Blue which was produced by autoclaving the 10% (v/v) Presto Blue solution. An increased fluorescence intensity indicated a greater reduction from resazurin to resorufin correlated with an increase in cell number. The % reduction of Presto Blue was calculated using equation (5).

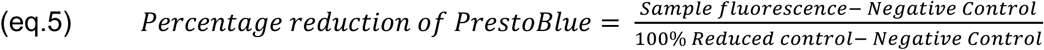

*(AlamarBlue Assay for Cell Proliferation | BMG LABTECH*, n.d.)

#### 2.6.3 Attachment efficiency assessment

Cell attachment was assessed at the 24 hour timepoint in order to allow cells time to attach and spread without proliferating. C2C12s were seeded at 2500 cells/cm². At 4 hours, wells were imaged using the incucyte S3 at 4 fields per well at the 10x magnification using phase contrast, confluency was analysed using the confluency analysis algorithm. Media was then removed cells were then washed twice in PBS and then fixed in 4% (v/v) paraformaldehyde for 10 minutes. Cells were then stained in a 300nM DAPI solution for 5 minutes before imaging on the Opera Phenix High throughput cell imager. Images of the whole well (96 well plate) were taken using the 5x air objective in non-confocal mode using the DAPI channel with 100 ms excitation, nuclei were then quantified using the count nuclei algorithm in the Columbus analysis software. Attachment score was then calculated using the below formula, assuming that the maximum expected confluency is 100% for satellite cells which has been observed previously.

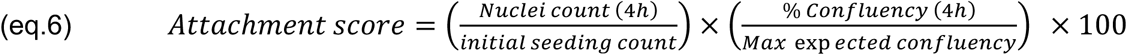

### 2.7 Immunocytochemistry

Cells were cultured in 48-well plates and fixed with 4% (v/v) paraformaldehyde for 20 minutes at room temperature, followed by two washes with PBS without Calcium and Magnesium. Samples were then permeabilised with 0.1% (v/v) Triton X-100 in 1× PBS for 15 minutes at room temperature and washed twice with PBS. Cells were then blocked with 5% (v/v) goat serum in 1× PBS for 45 minutes at room temperature in the dark. After removal of the blocking solution, they were incubated overnight at 4°C with primary anti-desmin antibody (mouse monoclonal, Sigma-Aldrich, SAB4200707) diluted to 1 µg/mL in PBS. Post-incubation, cells were subsequently washed three times with 1 mL/well cell staining buffer before incubation for 3 hours at room temperature in the dark with goat anti-mouse IgG1 cross-adsorbed secondary antibody conjugated to Alexa Fluor™ 488 (Sigma-Aldrich, A-21121), diluted to 2 µg/mL in PBS. Following secondary antibody incubation, wells were washed four times with 1 mL staining buffer per well (5 minutes per wash on a plate rocker). Nuclei were counterstained with 300 nM DAPI for 10 minutes at 37°C in the dark. After two final PBS washes, the cells were left in PBS and then imaged on the EVOS M5000 fluorescence microscope using FITC and DAPI filter sets.

Myogenic differentiation was quantified using a custom, open-source Python computer vision pipeline built upon the scikit-image and OpenCV libraries. To mitigate the variable background fluorescence characteristic of serum-free cultures, the myotube (Desmin) and nuclear (DAPI) channels were pre-processed using a White Top-Hat transform and Gaussian smoothing prior to intensity thresholding. Nuclei were segmented via a distance-transform watershed algorithm to separate touching cells, while myotube masks were refined using morphological area filters to exclude unspecific debris. Automated spatial analysis was then performed to calculate the strict fusion index, defined as the percentage of total nuclei residing specifically within mature myotubes (Desmin-positive regions containing ≥ 3 nuclei). The analysis script is publicly available at https://github.com/ucbewgo/Cell-Toolkit-Myo/.

### 2.8 Reverse Transcription Quantitative Polymerase Chain Reaction (RT-qPCR)

RNA extraction was performed using phenol/chloroform protocol. Briefly, cell pellets were disrupted in 1mL of TRIzol™ Reagent (Invitrogen). 200µL of chloroform was added and tubes were vigorously shaken. After 10min of incubation at RT, samples were centrifuged at 12,000g for 10min at 4°C. The aqueous phase was harvested and 500 µL of isopropanol was added. After 10min of incubation at RT, samples were centrifuged 12,000g for 8min at RT. Supernatant was carefully removed and 1mL of 75% ethanol was added to wash RNA pellet. Samples were centrifuged at 7,500 x g for 5 min and ethanol supernatant was removed. Pellets were air dried then dissolved in 50 µL of water then stored at -80°C.

500ng of RNA was harvested from each sample and treated with DNase using TURBO™ DNA-free kit (Invitrogen) following the manufacturer’s protocol. Then RNAs were reverse transcribed using the Superscript IV kit (ThermoFisher Scientific) following the manufacturer’s recommendation. cDNAs were diluted 5 times in water, and gene expression level was assessed by real-time quantitative PCR using the Power SYBR Green Master Mix (Thermo Fisher Scientific) on CFX Connect Real Time System (Biorad). Transcript RNA levels were normalized against GAPDH reference gene following the 2-ΔΔCt method. Full list of reagents. Primers can be found in the resource **T**able 2.

**Table 1:** List of qPCR reagents.

|  |  |  |
| --- | --- | --- |
| TRizol™ Reagent | Invitrogen | 15596026 |
| RNaseOUT | ThermoFischer | 10777019 |
| OligodT | ThermoFischer | 18418020 |
| dNTP | ThermoFischer | 18427013 |
| SuperScript IV Reverse Transcriptase | ThermoFischer | 18090050 |
| SYBR Green SsoAdvanced™ Universal Supermix | Biorad | 1725271 |
| TURBO DNA-free kit | Invitrogen | AM1907 |

**Table 2:**
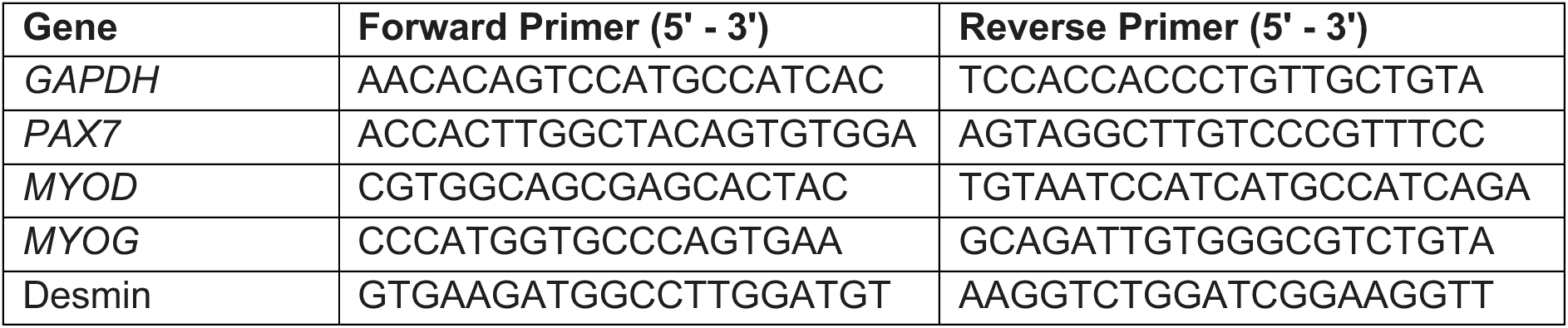
Primer Sequences.

### 2.9 Design of Experiments (DoE)

A sequential DoE approach was employed to systematically optimise the culture medium formulation. Initial screening of components was performed using Definitive Screening Designs (DSDs). Statistical analysis of these DSDs was conducted with forward stepwise regression and subset selection models. To control for model overfitting and the inflation of false positives across multiple factors, model selection was guided by the corrected Akaike Information Criterion (AICc) and Bayesian Information Criterion (BIC), rather than traditional independent null-hypothesis testing methodologies. This information-theoretic approach applies aggressive penalties for model complexity, ensuring that only statistically robust main effects and interactions are retained in highly saturated screening designs (B. Jones & Nachtsheim, 2011). Concurrently, residual analysis was used to incorporate quadratic terms where non-linear responses were evident. Subsequent optimisation rounds used full factorial and Central Composite Designs (CCDs) to investigate interaction effects and refine response surfaces. Models were iteratively fitted and assessed using R² and adjusted R², with non-significant terms systematically removed to enhance overall robustness and predictive power. All DoE screening matrices and statistical modelling were developed using JMP Pro versions 17 and 18.

### 2.10 Rheology

The dynamic viscosity of the MMM1 final formulation (with Growth medium (GM) as a control) with and without methylcellulose was measured at 25°C under shear stress ranging from 10—1000 s-1 using the Kinexus Lab + rheometer with a 40 mm measuring geometry (Malvern Instruments, Malvern, UK).

### 2.11 Cost Analysis

A laboratory-scale cost analysis was conducted to estimate and compare the raw material costs of the MMM1 formulation against standard serum-supplemented media (10% and 20% FBS) and previously published serum-free formulations (B8, Beefy-9, and Kolkmann et al., 2022). Cost estimations were calculated per litre (£/L) of prepared medium. Component pricing was sourced from standard commercial suppliers (e.g., Sigma-Aldrich, Thermo Fisher Scientific, R&D Systems) based on retail laboratory-scale quantities available via online pricing (e.g., milligrams for recombinant proteins, single litres for basal media and extracts) to reflect actual bench-scale procurement costs. To ensure an equitable biological and economic comparison, formulations heavily reliant on pharmaceutical-grade recombinant human serum albumin (hSA) were evaluated both by their full formulation cost and by their non-albumin cost.

### 2.12 Statistical Analysis

Statistical analysis and plotting was performed using Python (3.12.10) and GraphPad Prism (10.3.1). Differences between experimental conditions and the B8 control group were evaluated using Dunnett’s test for multiple comparisons. Data was expressed as mean ± standard deviation (SD). A minimum of three replicates were performed.

## 3. Results and Discussion

### 3.1 Establishing a baseline of growth and differentiation

The C2C12 cell line has been a preferred choice for many published studies focused on technologies for cultivated meat (Oh et al., 2023; O’Neill et al., 2023) due to its availability and its characteristics determined by their immortalised nature, as well as suitability for pet food applications. To benchmark the C2C12 cell line in a serum containing system and show it as a model for stable manufacturing of cultivated meat and pet food, its growth kinetics and differentiation potential in standard serum-containing conditions were first assessed and used as a baseline of comparison for serum-free media development with the aim to achieve similar or better cell growth than serum-based medium. C2C12 were cultured over 4 consecutive passages during which a non-significantly different doubling time of 20.92 ± 2.4 h (p > 0.05) was recorded during the 4 passages tested (**Figure 1A**). Additionally, C2C12 at passage 25 in serum-based medium were successfully differentiated towards the myogenic lineage as visualised by desmin staining of myotubes **(Figure 1B)** and phase contrast microscopy (**Figure 1C**).

**Figure 1:**
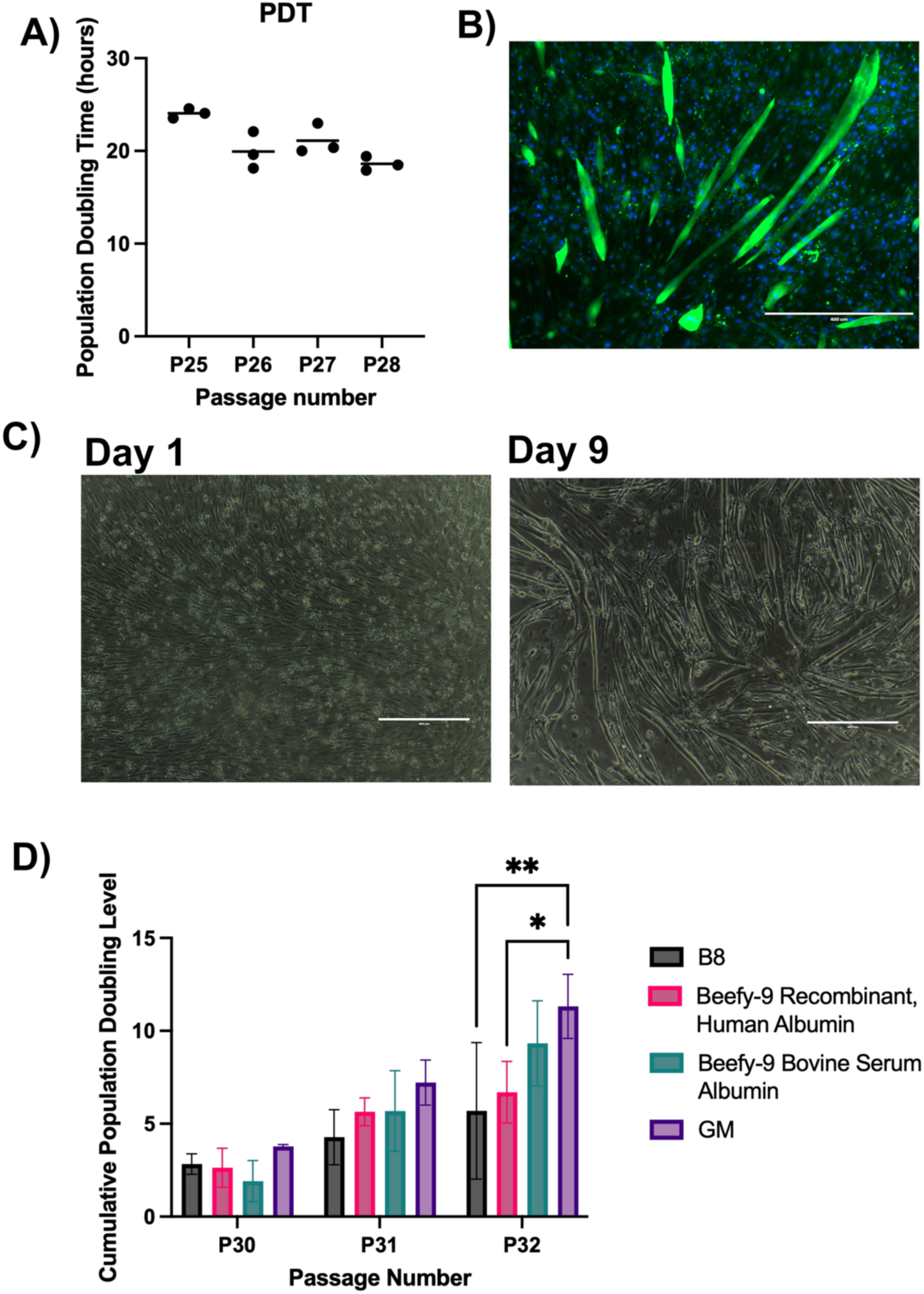
C2C12 baseline of growth and differentiation. (A) Population Doubling Time (PDT) (h) across four consecutive passages. Data shown as Mean ±SD (n=3); no significant difference was observed (One-way ANOVA). B) Representative fluorescence image of myotubes formed post-7 days of myogenic differentiation using serum starvation. Cells were stained with Desmin (green) for myotubes and DAPI (blue) for nuclei. Scale bar = 1000 µm. C) Phase contrast images of C2C12 cells at passage 25 differentiated towards the myogenic lineage for 9 days. Scale bar = 1000 µm. D) Cumulative Population Doubling Level (CPDL) of C2C12s cultured over three consecutive passages in B8 and Beefy-9 published serum-free media formulations compared to GM control. The serum albumin used in the Beefy-9 formulation was either bovine-derived or human recombinant. Data shown as mean ± SD (n=3). Significant differences were determined using a two-way ANOVA with Tukey’s post-hoc test (*p < 0.05, **p < 0.01).

Several serum-free media formulations have been published to date, one of which is Beefy-9 developed by Stout, Mirliani, et al. (2022) which is based on the B8 serum-free, animal-free media formulation initially developed for pluripotent stem cells (Kuo et al., 2020) with the additional supplementation of albumin. In this study, the B8 and Beefy-9 formulations were chosen as starting points. While both formulations were found to support C2C12 attachment and growth, a divergence in proliferation kinetics has emerged when tested over three consecutive passages. **Figure 1D** shows that by the 3^rd^ consecutive passage, the Cumulative Population Doubling Level (CPDL) of C2C12 murine myoblasts cultivated in B8 and Beefy-9 supplemented with recombinant human albumin were significantly lower than the serum-based control (GM) (*p < 0.05). These effects only became clear after multiple passages, highlighting the need to remove cells from serum for multiple passages to translate efficacy of serum-free media.

Albumin was found to be essential for supporting cell growth (Stout et al, 2022); however, it was also found to be one of the cost driving components of serum-free media (Kolkmann et al., 2020; Specht, 2020; Stout, Mirliani, et al., 2022; Stout, Rittenberg, et al., 2022). In this study, two different sources of albumin at different price points were tested – a cheaper bovine-derived variant (£0.003/mg) and a more expensive recombinant human variant (£1.03/mg). The source of albumin was found to have a significant effect on the performance of Beefy-9 formulation with C2C12 cells (*p < 0.05). The formulation supplemented with Bovine Serum Albumin (BSA) significantly outperformed the one containing Recombinant Human Albumin (**p < 0.01). This performance gap could be attributable to the “impurity” profile of animal-derived albumin or species-specific effects. Native albumin acts as a carrier protein that is naturally complexed with lipids, fatty acids, and trace micronutrients *in vivo* (Francis, 2010). In contrast, pharmaceutical-grade recombinant albumin is highly purified and potentially stripped of these associated ligands (Maier et al., 2021; Petitpas et al., 2001). In this particular study, the poorer performance of the recombinant variant with C2C12 murine myoblasts could be linked to the absence of lipids and fatty acids typically carried by serum albumin, which make recombinant albumin act as a “nutrient-sink” through the high affinity binding (Ferrer et al., 2001) to other recombinant proteins present in the media formulation (Keenan et al., 2006; Larsen et al., 2016). Further testing would be required to verify this hypothesis and to achieve a better understanding of the effects seen. However, the purpose of this manuscript was not to investigate the mechanisms behind the different components, but to design and develop a serum-free media formulation that performs equally if not better than serum-based media. From a cost perspective, the animal-derived albumin variant is also cheaper than the recombinant one making it attractive for cultivated meat applications; however, it is an animal-derived component which can introduce additional challenges such as batch-to-batch variability and impurities (Boelt et al., 2022).

### 3.2 Identification of the Key Media Components and their effect on C2C12 cell growth

To understand the key components in the B8 and Beefy-9 formulations that have an effect on C2C12 cell proliferation, a Definitive Screening Design of Experiments (DoE) was performed. The component concentrations were either kept as they are in the original published formulations (Stout et al, 2022), halved or completely removed. This approach facilitated the screening of both main effects and second-order (quadratic) interactions by systematically reducing component concentrations. Full experimental conditions can be found in the **Supplementary Table 1**.

The screening revealed a significant positive quadratic effect for FGF2 (FGF2*FGF2, *p < 0.05) suggesting that the relationship between FGF2 concentration and cell proliferation is a non-linear progression (**Supplementary Figure S1**). Nuclei count **(Figure 2A)** showed a drop in cell growth at 20 ng/mL FGF2 compared to 0 or 40 ng/mL suggesting that the optimal concentration lies within a specific range, and deviations from this optimum (either too low or saturation levels) yield diminishing returns. Another significant effect was seen for albumin which exerted a significant negative effect on the C2C12 proliferation (*p < 0.05). It is worth mentioning that the recombinant human variant was used in this screen. This inhibitory profile was observed across both nuclei enumeration **(Figure 2A)** and metabolic activity **(Figure 2B)**, suggesting that the removal or reduction of recombinant albumin in this formulation is critical for achieving C2C12 murine myoblast growth.

**Figure 2:**
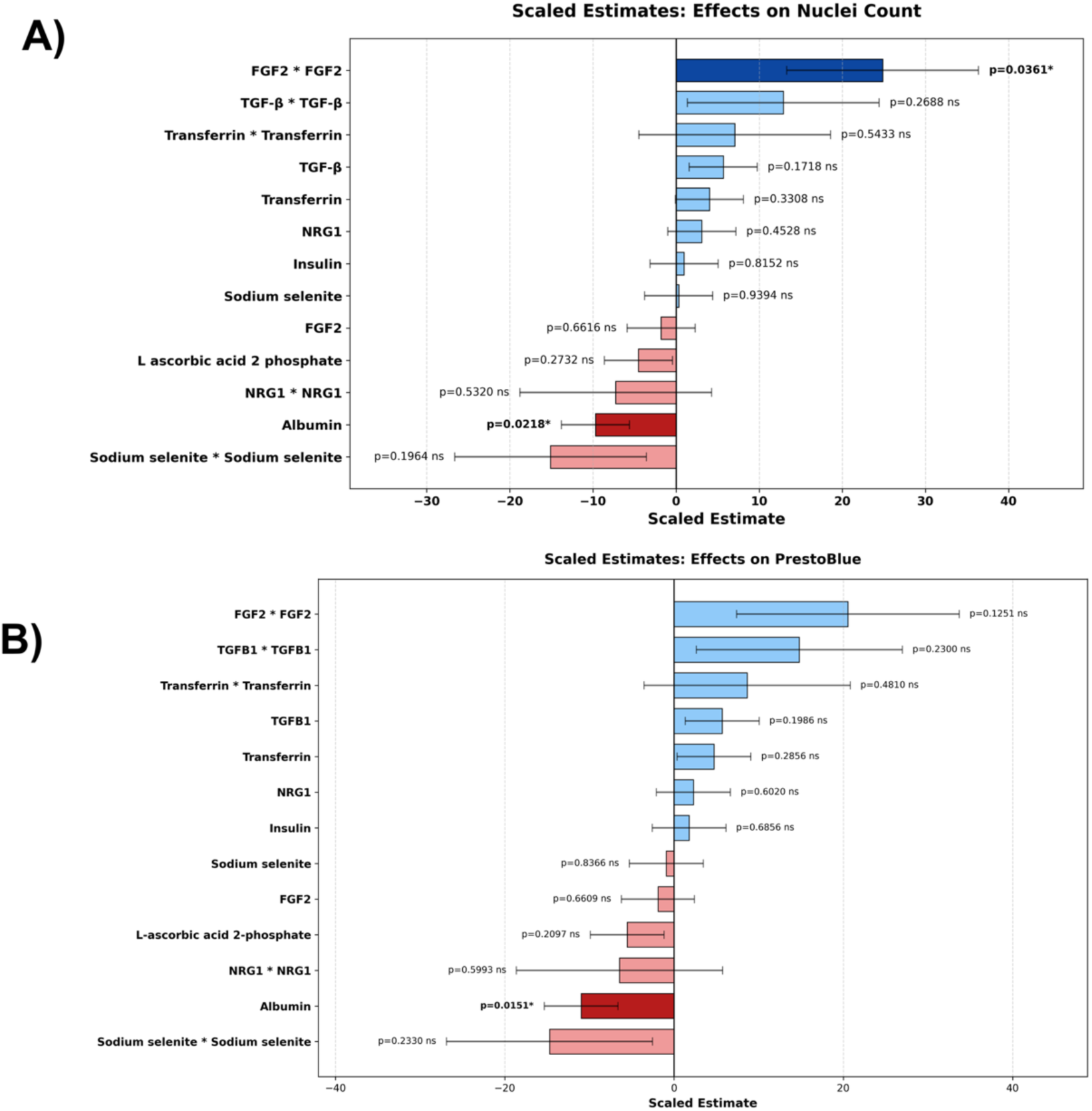
Identification of key media components in the published B8 and Beefy-9 formulations (Stout et al, 2022) and their effects on C2C12 murine myoblasts proliferation. (A–B) Scaled estimates from the Definitive Screening Design (DSD) identifying the main and quadratic effects of media components on (A) Nuclei Count and (B) Metabolic Activity (Presto Blue assay). Bars represent the standardised effect size; blue indicates a positive influence on proliferation, while red indicates a negative influence. P-values are annotated adjacent to each term. Albumin (Human Recombinant) showed a significant negative effect (p < 0.05), FGF2 quadratic effect showed a significant positive effect (p < 0.05).

Additional growth factors with potential to support cell growth were also investigated in a one-factor-at-a time (OFAT) approach and these included IGF-1, HGF, IL-6, VEGF, and PDGF-BB added to a basal medium of B8 **(Figure 3A).** These growth factors were chosen as they have been found to promote cell proliferation (Cantini et al., 1995; Machida & Booth, 2004; Suzuki et al., 2000) and extracellular matrix (ECM) deposition in satellite cells (Albrecht & Tidball, 1997). Most growth factors tested elicited similar or slightly higher fold change relative to the B8 control. However, only one factor produced a statistically significant increase in C2C12 proliferation (*p < 0.05) and that was HGF, but only at the lowest concentration tested of 0.25 ng/mL. HGF is known to play a role in activating quiescent satellite cells and promoting early-stage myoblast division (Sheehan et al., 2000). However, high concentrations of HGF have been found to be inhibitory to myoblast proliferation (Yamada et al., 2009) which was also seen here with concentrations above 0.5 ng/mL (**Figure 3A**).

**Figure 3:**
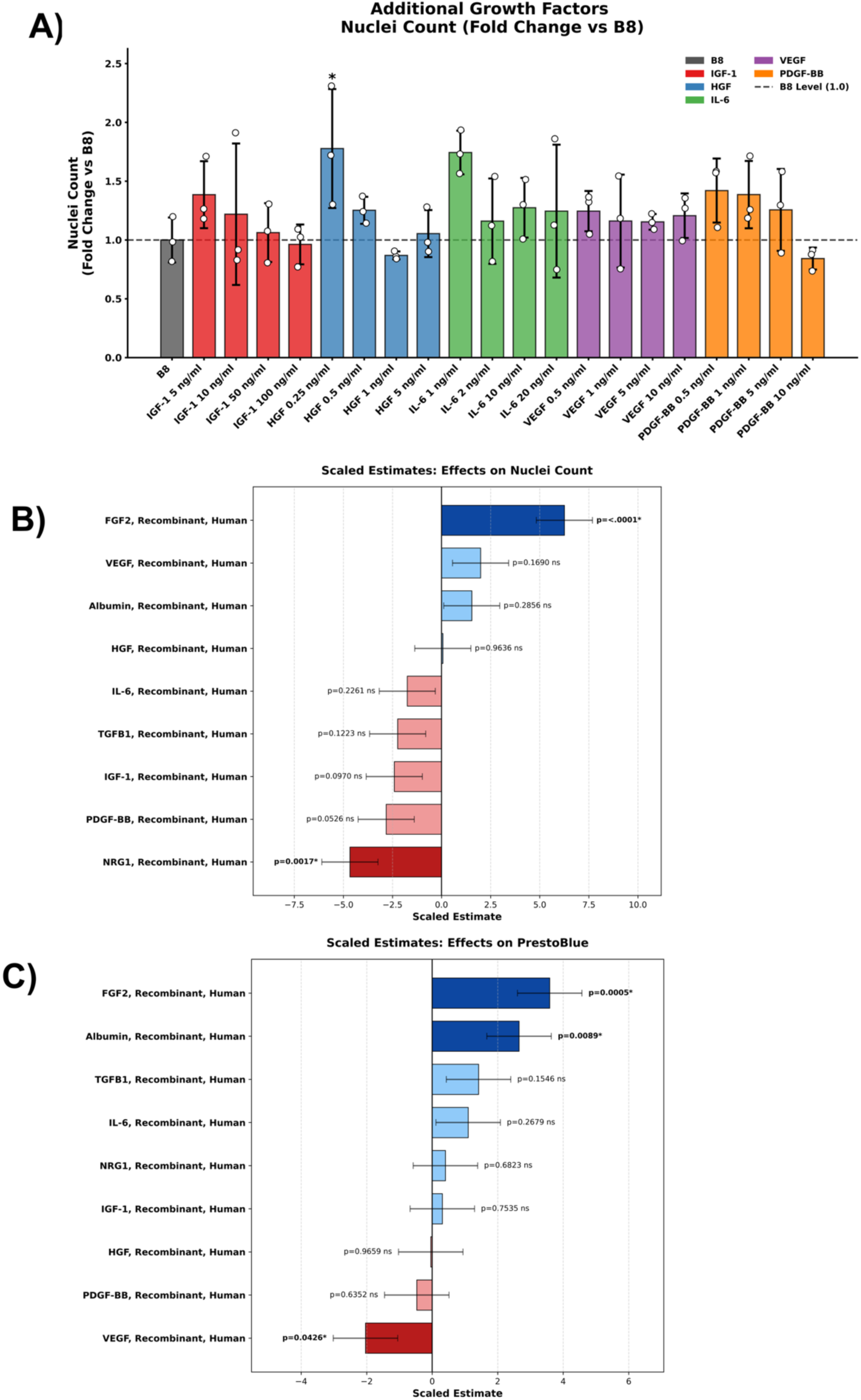
A) Dose-response screening of additional growth factors (IGF-1, HGF, IL-6, VEGF, and PDGF-BB) supplemented to the basal B8 medium. Data are presented as fold change relative to the B8 negative control (set to 1.0, dashed line). Data shown as mean ± SD (n=3). Individual data points indicate independent experimental values. Statistical significance was determined using independent Dunnett’s Post Hoc Test. Asterisks denote significance relative to B8 control (*p < 0.05). (B) Scaled estimates from the Definitive Screening Design (DSD) on Nuclei Count. Horizontal bars represent the standardised effect size of individual components. Blue bars indicate a positive influence on cell number, and red bars indicate a negative influence. Specific significance is noted with p-values, highlighting a significant negative impact of NRG1 and a strong positive impact of FGF2 (p < 0.05). (C) Scaled estimates from the DSD on Metabolic Activity (PrestoBlue assay). Results follow a similar pattern, where horizontal bars display standardized effect size. VEGF shows a significant negative influence (red bar, p < 0.05), while Albumin and FGF2 have significant positive effects (blue bars, p < 0.05) on overall metabolic activity. For clarity, P-values are annotated adjacent to each factor term.

To move beyond one-factor-at-a-time (OFAT), a definitive screening design (**Supplementary Table 2**) was adopted to test the combination of these growth factors in a combined media formulation. In this screen, FGF2, once again had a significant positive effect on both nuclei count (**Figure 3B**) and Presto Blue (**Figure 3C**), confirming what was found in the previous DoE and OFAT screens that FGF2 is a key driver of C2C12 proliferation. Contrary to the effects seen in **Figures 2A & 2B** where albumin had a negative effect, in this screen albumin had a significantly positive effect on C2C12 metabolic activity as measured by Presto Blue. This may be that the positive effect of albumin as a protein carrier, whereby it binds and carries proteins into cells (Larsen et al., 2016; Schenzle et al., 2025) is exacerbated with additional growth factor proteins added to the mixture. This model did not, however, identify significant interaction effects between albumin and other growth factors. A more detailed DoE design such as a central composite design (Almeida et al., 2025) may be able to identify such interaction effects.

### 3.3 Screening of Food-grade Additives and Macro-Molecular Crowders

Following the identification of recombinant albumin as a component impacting cell growth negatively, alternatives to albumin from low-cost, food-grade sources were also investigated. A panel of candidates including methylcellulose, beta-carotene, curcumin, trehalose, racemic alanine, and L-carnosine added to a basal medium of B8 was screened via a OFAT approach **(Figure 4A).** The selection of these specific components was done as to replace the different functions that albumin typically carries out such as protein binding/carrying (e.g. trehalose) (Olsson et al., 2016), antioxidant activity (e.g. beta-carotene, curcumin, L-carnosine) (Ak & Gülçin, 2008; Liu et al., 2022; Tsuchihashi et al., 1995) and the control of the physical properties of the media (e.g. methylcellulose) (Schenzle et al., 2025).

**Figure 4:**
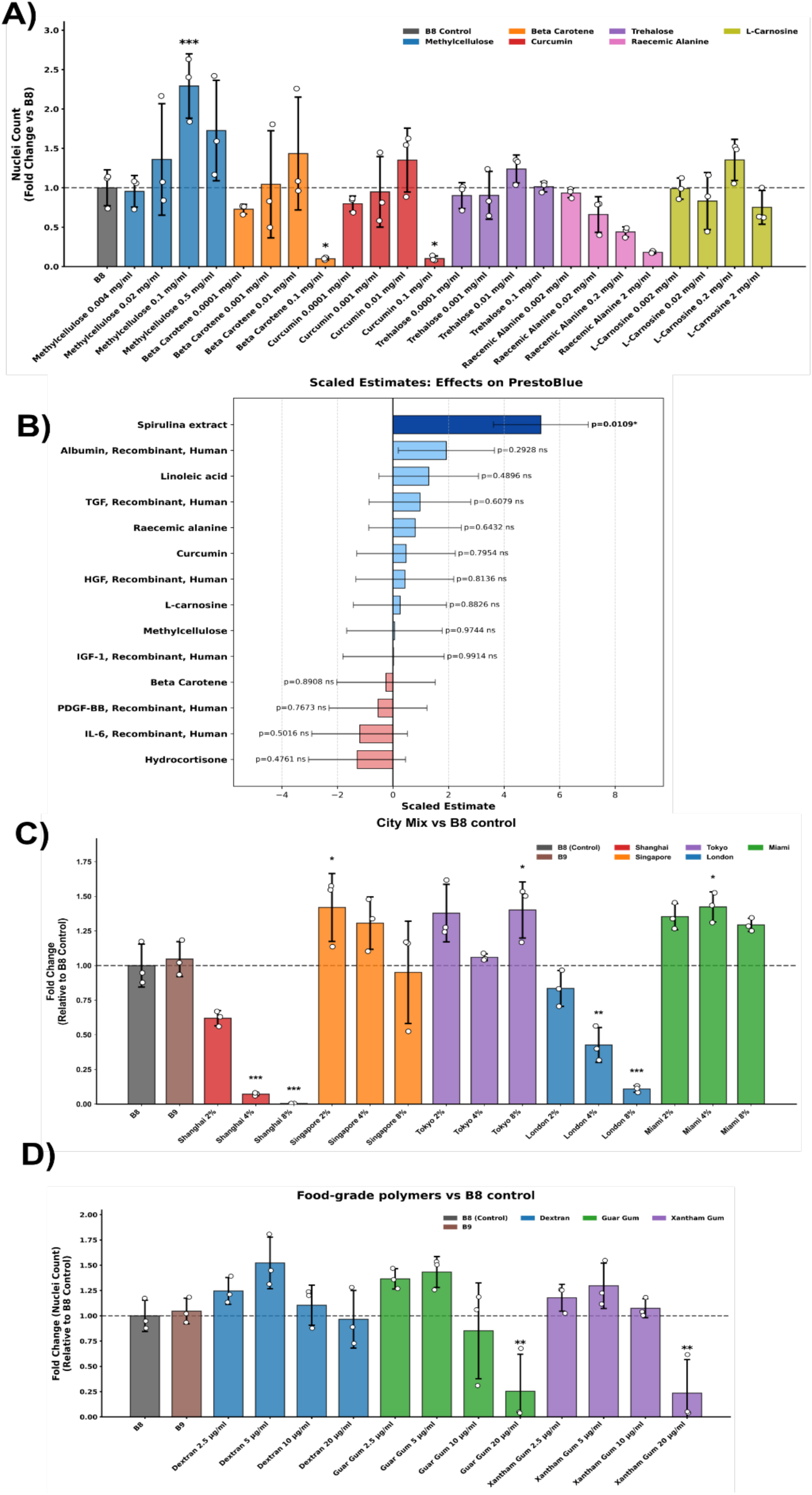
Screening of food-grade candidates and macromolecular crowders for albumin replacement. A) Dose-response screening of selected food-grade additives (Methylcellulose, Beta Carotene, Curcumin, Trehalose, Racemic Alanine, and L-Carnosine) on C2C12 nuclei count. Data is expressed as fold change relative to the B8 control (dashed line). A significant increase was observed for Methylcellulose (0.1 mg/mL). B) Scaled estimates from a design of experiments (DoE) screen identifying the effects of various food-grade components on metabolic activity (Presto Blue). Spirulina extract showed a significant positive effect (p = 0.0109). C) Efficacy of commercial “City-Mix” macromolecular crowding supplements (Shanghai, Singapore, Tokyo, London, Miami) at varying concentrations. Data is shown as mean ± SD (n=3). Significant differences compared to the B8 control were determined using a one-way ANOVA with Dunnett’s multiple comparisons test (*p < 0.05, **p < 0.01, ***p < 0.001). D) Evaluation of food-grade polymers (Dextran, Guar Gum, Xanthan Gum) as potential macromolecular crowders. Nuclei counts are normalised to the B8 control. Data is shown as mean ± SD (n=3).

Amongst the candidates and their range of concentrations tested, methylcellulose at a concentration of 0.1 mg/mL was the only additive that resulted in a statistically significant C2C12 growth increase relative to the B8 control (*p < 0.05). Methylcellulose is a chemically inert polymer often used to modify medium viscosity. In this study, the physical modulation of the culture environment generated by methylcellulose proved to be a critical driver of C2C12 growth. On the contrary, supplements like L-carnosine and trehalose didn’t result in cell growth at any of the tested concentrations. Conversely, lipophilic antioxidants such as curcumin and beta-carotene proved cytotoxic at the higher concentrations tested, significantly reducing the C2C12 murine myoblast growth (*p < 0.05), likely as a result of oxidative stress or membrane disruption. At the lower concentrations of beta-carotene and curcumin, no such significant negative effects were observed, therefore these components might have other utility as potential antioxidants and that was not reflected in the nuclei count in this case (Ak & Gülçin, 2008; Tsuchihashi et al., 1995). Racemic alanine has previously been found to have a stabilising effect on growth factors (Benington et al., 2021; Schenzle et al., 2025). However, here it was found to inhibit C2C12 growth in the concentration range tested with an inhibitory response proportional to the increase in concentration. While L-carnosine was found to have positive effects on porcine satellite cell proliferation (Liu et al., 2022), in this study, no significant difference in C2C12 cell growth was found. This may be due to species-specific effects or an effect of serum presence in the Liu et al (2022) study unlike this study with serum-free conditions.

To further understand the effects of bioactive components in complex media formulations on C2C12 murine myoblast growth, another Definitive Screening DoE was conducted. Full design can be seen in **Supplementary Table 3**. This screen incorporated a mix of growth factors with potential (e.g. HGF, IGF, TGFβ, PDGF-BB, IL-6), and food grade components (e.g. spirulina extract, racemic alanine, methylcellulose, linoleic acid, curcumin, beta-carotene) (**Figure 4B**). Spirulina extract has been selected as it was found to support murine myoblast growth in other studies (Gordon-Petrovskii et al., 2026). In this screen, only the spirulina extract produced a significant positive effect on metabolic activity (*p = 0.0109 < 0.05) of the C2C12 murine myoblasts. This significant efficacy likely stems from a multi-modal mechanism of action rather than a single active compound. Spirulina is a nutrient-dense cyanobacterium containing phycocyanin, complex proteins, vitamins, and essential fatty acids (Choopani et al., 2016; Clément et al., 1967; Liestianty et al., 2019; Stunda-Zujeva et al., 2023). A concurrent compositional analysis of the exact extract used in this study (Gordon-Petrovskii et al., 2026) revealed it is particularly high in proteins, while remaining relatively low in free amino acids. In a serum-free environment, this complex profile allows the extract to act simultaneously as a potent antioxidant (mitigating oxidative stress via phycocyanin), a source of bioavailable lipids, and a provider of trace micronutrients that are traditionally supplied by serum but absent in chemically defined basal media. While biophysical fractionation would be required to fully map these individual lipidomic and proteomic interactions, the use of the whole extract provides a highly functional, food-grade solution to the nutritional gaps of defined media. No significant interaction effects were observed in this screen.

To assess the robustness of this food-grade supplementation, we evaluated the performance of three distinct commercial spirulina extract batches (Batch 1, Batch 2, from the same manufacturer - Naturya and an additional batch from a different manufacturer - Bulk Powders). As shown in **Supplementary Figure S2**, batch-to-batch variability was evident. While there was no significant difference between the Naturya batch 1 (same as used in all development studies) and the Bulk Powders source, a significant decrease in cell yield (p < 0.05) was observed when comparing the Naturya batch 2 to batch 1. This difference in cell performance could be attributed not only to batch-to-batch variation, but also to shelf life as batch 1 was stored for a period of 18 months at room temperature prior to these experimental runs, while batch 2 was used within 5 days of purchase.

Methylcellulose was found to have a positive effect on C2C12 murine myoblasts cell growth, likely through its established ability to increase media viscosity and provide structural hydrodynamic shear protection (Goldblum et al., 1990; Schenzle et al., 2025). To explore whether physical modulation of the culture environment could be further enhanced through true macromolecular crowding, a phenomenon where inert molecules mimic the dense *in vivo* environment by occupying volume (Zhou et al., 2008), five proprietary crowding supplements from the ‘City-Mix’ range (3D BioTissues, UK) were evaluated via OFAT addition to a B8 basal medium (**Figure 4C**). The poorest performing City-Mix tested was Shanghai-Mix which was found cytotoxic with significant reduction in C2C12 growth particularly at 4% (v/v) and 8% (v/v) (***p < 0.001). London-Mix behaved similarly with significant reduction in C2C12 growth at the higher concentrations (**p < 0.01 and ***p < 0.001). On the opposite, the Singapore-Mix at 2% (v/v), the Tokyo-Mix at 8% (v/v), and the Miami-Mix at 4% (v/v) significantly increased the C2C12 murine myoblasts growth compared to the B8 control (*p < 0.05). However, the exact formulations of the City-Mixes are proprietary, and as such we wanted to determine if these crowding effects could also be replicated using generic food-grade polymers that have been previously reported to act as macromolecular crowders. As such, dextran, xanthan gum, and guar gum were selected based on previous reports (Guillaumin et al., 2024; Lareu et al., 2007; Prewitz et al., 2015) and tested **(Figure 4D)**. While minor, non-significant increases in nuclei counts were observed at the lower concentrations, these generic polymers failed to recapitulate the significant efficacy obtained with the City-Mix formulations. Furthermore, higher concentrations of guar gum and xanthan gum were found to be inhibitory to C2C12 cell growth, likely due to increased viscosity impeding the diffusion of nutrients and waste products (Ahmad et al., 2022).

### 3.4 An effective serum-alternative formulation developed and validated

The Definitive Screening DoE described in **Figure 4B** was also used to suggest an optimal formulation based on its results. The JMP software provides an “optimisation profiler” for the DoE model which allows a model predicted optimal formulation. Using this tool, an optimal formulation candidate named here the “*Mus Musculus* Medium 1” (MMM1) (shown in **Figure 5A**) was further selected for validation studies.

**Figure 5:**
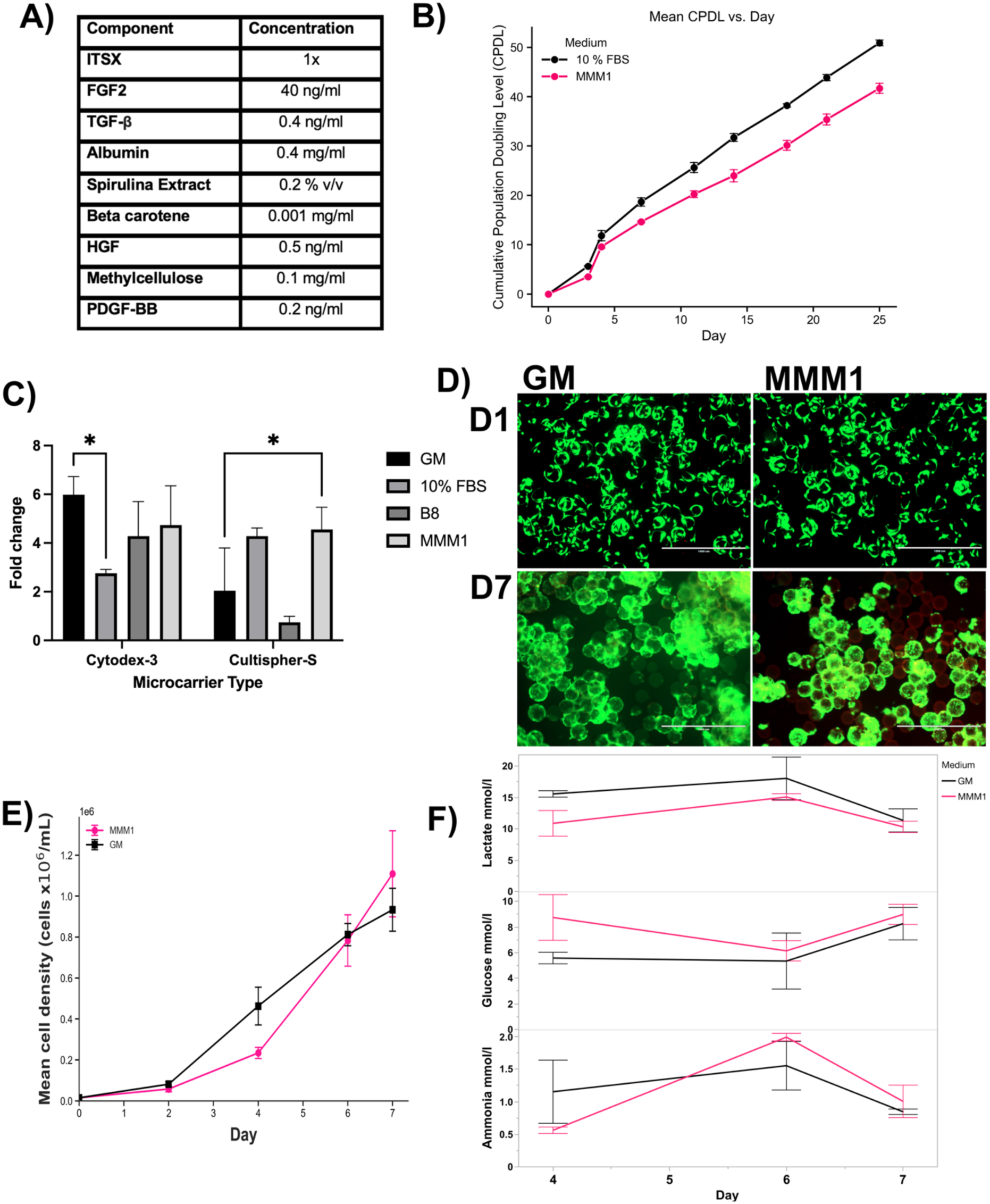
Validation of MMM1 serum-free medium formulation in C2C12 murine myoblast culture. (A) MMM1 Formulation (B) Cumulative Population Doubling Level (CPDL) of C2C12 cells cultured in the MMM1 optimised serum-free formulation compared to the standard serum control (10% (v/v) FBS) over 29 days of culture in microwell plates seeded at 2,500 cells/cm^2^. Data shown as mean ± SD (n=3). Expansion of C2C12 myoblasts on (C) Cultispher-S and Cytodex-3 microcarriers in MMM1 serum-free media formulation in static conditions. Data shown as mean ± SD (n=3). (D) Representative Live/Dead stain images of C2C12 murine myoblasts cultured on Cultispher-S microcarriers in MMM1 serum-free formulation or GM serum-based formulation in agitated cultures in spinner flasks. Images taken at Day 1 and Day 7 of cultures. Live cells are stained green (Calcein AM), while dead cells are stained red (Ethidium Homodimer). Scale bars = 1000 µm. (E) Cell densities of C2C12 murine myoblasts cultured on Cultispher-S in spinner flasks over 7 days. By Day 7 compared to the GM control. Data shown as mean ± SD n=5. Crucially, n=5 represents three entirely independent spinner flask runs, with each replicate executed in a separate vessel using a distinct cell inoculum of C2C12s at P23, P24 and P27. Each spinner flask run consists of two spinner flasks. (F) Metabolic profiling and spent media analysis of C2C12 myoblasts during microcarrier expansion. Concentrations of ammonia, glucose, and lactate were quantified over a 7-day culture period in 100 mL agitated spinner flasks. Cells were cultivated on Cytodex-3 microcarriers in either the custom serum-free formulation (MMM1, blue line) or the standard 20% serum-containing control (GM, red line). A semi-continuous feeding regime was implemented, consisting of an 80% media volume exchange on days 2, 4, and 6. Data points represent spent media samples extracted immediately prior to these feeding interventions. Data are presented as mean ± SD (n = 3).

To clearly delineate the novelty of the MMM1 formulation, a side-by-side component comparison against established, highly-cited serum-free satellite cell expansion media including B8/Beefy-9 (Kuo et al., 2020; Stout, Mirliani, et al., 2022; Kolkmann et al., 2022), and (Skrivergaard et al., 2023) is provided in **Supplementary Table 4**. While MMM1 shares foundational proliferative drivers (such as FGF2, TGF-beta, and ITS) with elements of these formulations, its distinct novelty lies in lower concentrations of carrier proteins such as albumin or fetuin. Instead, MMM1 successfully introduces unique, lower-cost components, leveraging alternatives that can be food grade, specifically spirulina extract, methylcellulose, and beta-carotene.

In a first instance, the MMM1 formulation was evaluated in monolayer culture over eight consecutive passages and compared to serum-based GM. The MMM1 serum-free medium formulation achieved a Cumulative Population Doubling Level (CPDL) of 41.7 ± 1, lower than the CPDL of 50.9 ± 0.56 achieved in the serum-containing control (10% FBS) over 8 consecutive passages equivalent of 29 days in culture (**Figure 5B**). Differentiation potential was maintained after 4 passages in both serum-containing and serum-free medium (**Supplementary Figures S3 & S4**) with a mean fusion index of 21.9% (from a mean of 1008.67 nuclei per image) vs GM 10.9% (from a mean of 965 nuclei per image) from an (p=0.27). Furthermore, RT-qPCR analysis was conducted on samples cultured in the long-term study shown in **Figure 5B**. The analysis confirmed differentiation (based on relative RNA levels of MYOD, MYOG and Desmin) of samples cultured in 10% FBS and MMM1 after 3 and 7 passages in the respective media (**Supplementary Figure 5**).

The next step was to validate the suitability of the MMM1 serum-free medium formulation for scalable bioprocesses. As C2C12 murine myoblasts are adherent dependent, the MMM1 formulation was tested in microcarrier cultures first in static conditions in ultra-low attachment well plates (**Figures 5C–D**) and then agitated conditions in 100 mL spinner flasks (**Figure 5E**). Two commercially available microcarriers with different surface chemistries and properties were chosen. These were Cultispher-S (Percell Biolytica) which is macroporous, gelatin-based microcarrier and Cytodex-3 which is a microporous, dextran-based microcarrier coated with denatured collagen (de Bournonville et al., 2021). On both microcarriers tested, the C2C12 murine myoblasts cultured in the MMM1 serum-free media formulation successfully attached and grew over time as seen by live/dead staining (**Figure 5D**). By day 7 in static conditions, microcarrier aggregates were formed indicating cell proliferation over time as was previously seen with microcarrier cultures (Hanga et al., 2020, 2021). In 10 days of static culture, a fold change of 4.73 ± 1.6 h was achieved in MMM1 on Cytodex-3 compared to 4.55 ± 0.9 h on Cultispher-S (**Figure 5C**). When C2C12 were cultured on Cultispher-S, a significantly higher fold change was achieved in MMM1 compared to growth medium containing 20% (v/v/) serum (*p < 0.05), while on Cytodex-3, no significant difference in cell growth was achieved between GM and MMM1 formulation (**Figure 5C**). These differences in cell growth on Cytodex-3 compared to Cultispher-S microcarriers are closely linked to the surface chemistry of these microcarriers which elicit different cell responses from attachment to proliferation rates (Andreassen et al., 2022).

Following static culture validation, the MMM1 formulation was then validated in dynamic culture in 100 mL spinner flasks. Cell attachment and viability on microcarriers was confirmed with Live/Dead staining. For this validation, only Cultispher-S microcarriers were selected based on their growth comparable to the growth medium (GM) control. The C2C12 murine myoblasts had similar growth profiles in the MMM1 serum-free formulation compared to the serum-containing GM on Cultispher-S microcarriers with overpassing serum-based medium by day 7 of culture when an average final cell concentration of 1.11 x 10^6^ cells/mL was achieved in the MMM1 serum-free formulation compared to 0.933 x 10^6^ cells/mL in the GM condition (**Figure 5E**). These findings demonstrate that the MMM1 serum-free medium formulation is able to support the growth of C2C12 murine myoblasts even in scalable bioreactor systems while achieving comparable growth to serum-based medium. Future pilot-scale studies must evaluate enzymatic detachment kinetics to ensure high harvesting viability.

To evaluate the metabolic sustainability of the MMM1 formulation during high-density suspension culture, preliminary spent media analysis was conducted on days 4 and 7 of a representative spinner flask batch (using an 80% media exchange regime on days 2, 4, and 6) (**Figure 5F**). Tracking macronutrient fluxes revealed an active metabolic profile that accelerated alongside cellular expansion. By Day 7, glucose levels in the MMM1 culture dropped to an average of 8.99 mmol/L, reflecting rapid carbon metabolism. Correspondingly, lactate accumulation reached 11.50 mmol/L in MMM1, which notably remained below the serum-containing control (14.59 mmol/L), indicating that the MMM1 formulation did not exacerbate anaerobic glycolytic flux. Ammonia accumulation was comparable, reaching a maximum of 1.14 mmol/L in MMM1 versus 0.90 mmol/L in the control. Overall, these trajectories confirm that MMM1 effectively supports high-density metabolic demands, though future pilot-scale transitions will require optimised, continuous perfusion feeding to prevent late-stage glucose exhaustion.

To further evaluate the biophysical properties of the final MMM1 formulation, its steady-state shear viscosity was measured (**Supplementary Figure S5**). The baseline medium lacking methylcellulose (MMM1 - MC) and the serum-supplemented control (GM) exhibited relatively flat viscosity profiles at low-to-moderate shear rates. In contrast, the addition of methylcellulose (MMM1 + MC) imparted a distinct non-Newtonian, shear-thinning profile. At low shear rates (10 s⁻¹), the supplemented medium maintained an elevated viscosity (approximately 1.5 × 10⁻¹ Pa·s). Conversely, at high shear rates (10³ s⁻¹), the viscosity droped by over an order of magnitude, converging with the base fluids. This shear-responsive behaviour, which maintains a higher viscosity in low-shear zones while thinning under higher shear stresses, is consistent with the rheological properties typically required to support and protect cells during dynamic agitated culture.

### 3.5 Further optimisation of the MMM1 serum-free medium formulation for Animal-Free Compliance and Cost Reduction

The initial iteration of the MMM1 serum-free formulation relied on the use of commercial insulin-transferrin-selenium-ethanolamine (ITS-X) and bovine serum albumin to stabilise its performance, however both components are animal-derived. Animal-free formulations are highly desirable as they are more compliant with regulations. As such, to achieve a fully defined, animal-free formulation, a final optimisation screen was conducted using a DoE approach (full design in **Supplementary Table 5**) to substitute the commercial ITS with an “Optimal ITS” blend derived from the previous screening phase (2500 ng/mL insulin, 4591 ng/mL transferrin, 10.05 ng/mL sodium selenite), as well as an animal-free commercial variant. Additionally, recombinant human albumin was re-evaluated to determine if the improved nutritional context of the MMM1 formulation containing spirulina and methylcellulose could restore its utility.

In contrast to the inhibitory effects observed in the unoptimised B8 basal medium, recombinant human albumin in this improved MMM1 formulation demonstrated a significant positive effect on the metabolic activity of the C2C12 murine myoblasts (**Figure 6B**; *p=0.0048 < 0.05). This new result suggested that the inclusion of food-grade supplements such as spirulina extract and methylcellulose, through their stimulation of metabolic and proliferation rates were able to counterbalance any negative effects of the recombinant albumin seen in the previous screen. Furthermore, the custom “Optimal ITS” formulation significantly enhanced the nuclei count (*p=0.0205 < 0.05) (**Figure 6A**) compared to the standard animal-free alternative, indicating that optimising the stoichiometric ratios of insulin and transferrin is more critical than the source origin alone. High concentrations of FGF2 (40 ng/mL) and TGF-β (0.4 ng/mL) were also confirmed as essential drivers of proliferation. Consequently, the final animal-free, serum-free MMM1 formulation was defined to include Recombinant Albumin, the “Optimal ITS” blend, and sustained levels of FGF2 and TGF-β.

**Figure 6:**
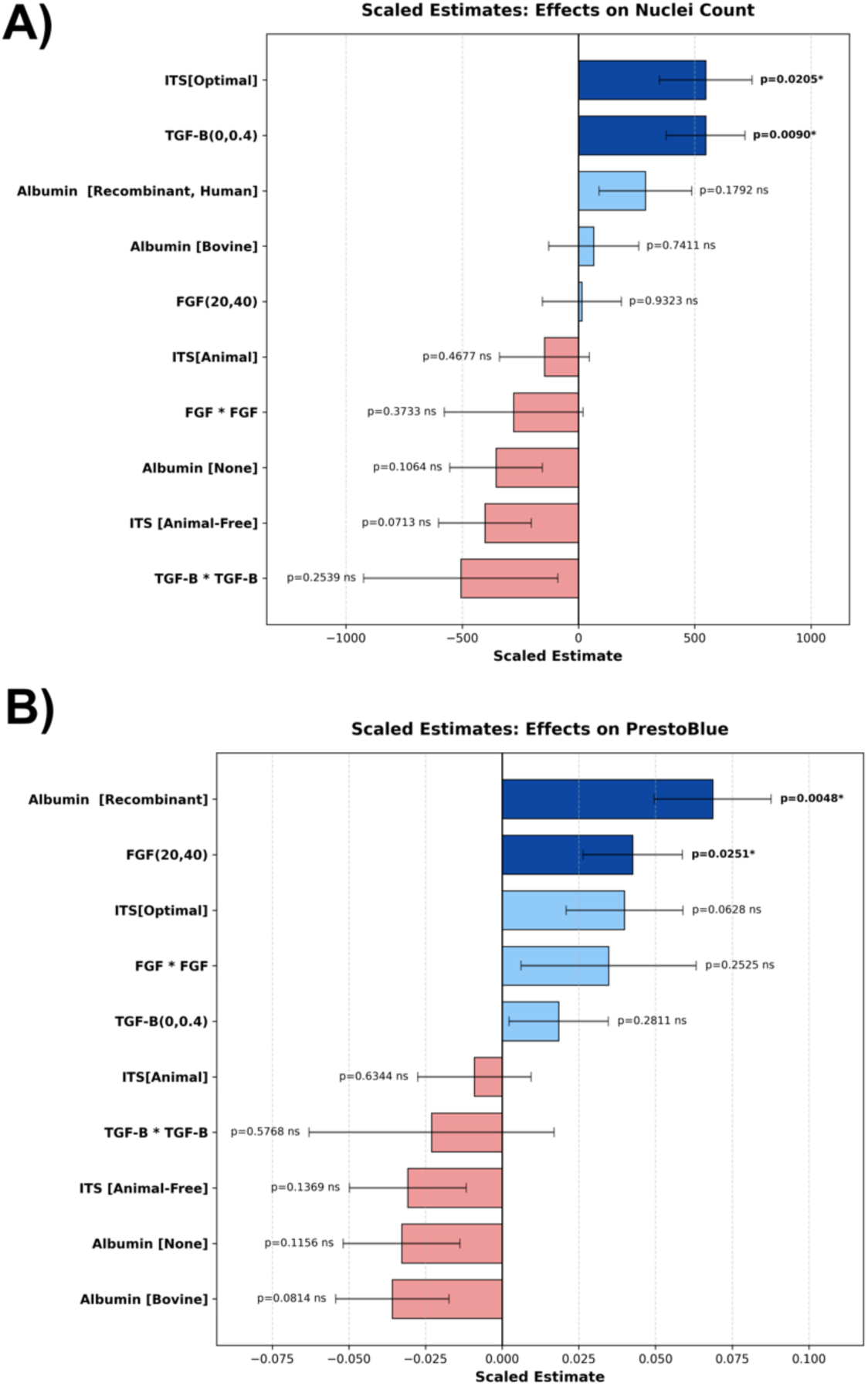
Further optimisation of the MMM1 serum-free formulation for cost reduction and elimination of animal-derived components. (A) Scaled estimates from a design of experiments (DoE) screen evaluating the impact of component substitutions on Nuclei Count. The “ITS [Optimal]” formulation and TGF-β exhibited significant positive effects on proliferation (p < 0.05). (B) Scaled estimates of component effects on metabolic activity (Presto Blue). Recombinant Albumin and FGF2 (20/40 ng/mL) significantly increased metabolic activity (p < 0.05). Data shown as mean ± SD (n=3).

Beyond protein supplements, the basal medium itself represents a dominant cost driver in large-scale bioprocesses due to the sheer volumes required. To address this, the performance of the C2C12 murine myoblasts was compared when cultured in standard Pharmaceutical-Grade DMEM/F12 (Gibco, Thermofisher) versus a bulk Food-Grade DMEM/F12 variant (Multus Biotechnology). Remarkably, the transition to food-grade basal media resulted in no statistically significant difference in proliferation of C2C12 murine myoblasts over a 10-day culture period (**Figure 7A**; p > 0.05). Similarly, the C2C12 attachment efficiency on uncoated surfaces remained comparable between the two grades of basal medium tested (**Figure 7B**). These findings confirmed that the strict purity standards required for pharmaceutical applications are not biological prerequisites for muscle cell expansion. The successful validation of food-grade basal media provides a promising strategy to substantially reduce media costs in cultivated meat production without compromising biological performance. The price of pharma grade DMEM/F12 is between £48-70/L whereas the food-grade DMEM/F12 tested currently costs £7.5/L with lower prices in bulk powder format. Furthermore, it is estimated that at food grade basal media can cost as little as £0.15/L with costs from amino acids contributing as little as £0.015-0.13/L when sourced from food or feed-grade sources (Gibbons et al., 2025). This therefore demonstrates the efficacy of food-grade basal media and its potential for further lowering cultivated meat input costs.

**Figure 7.**
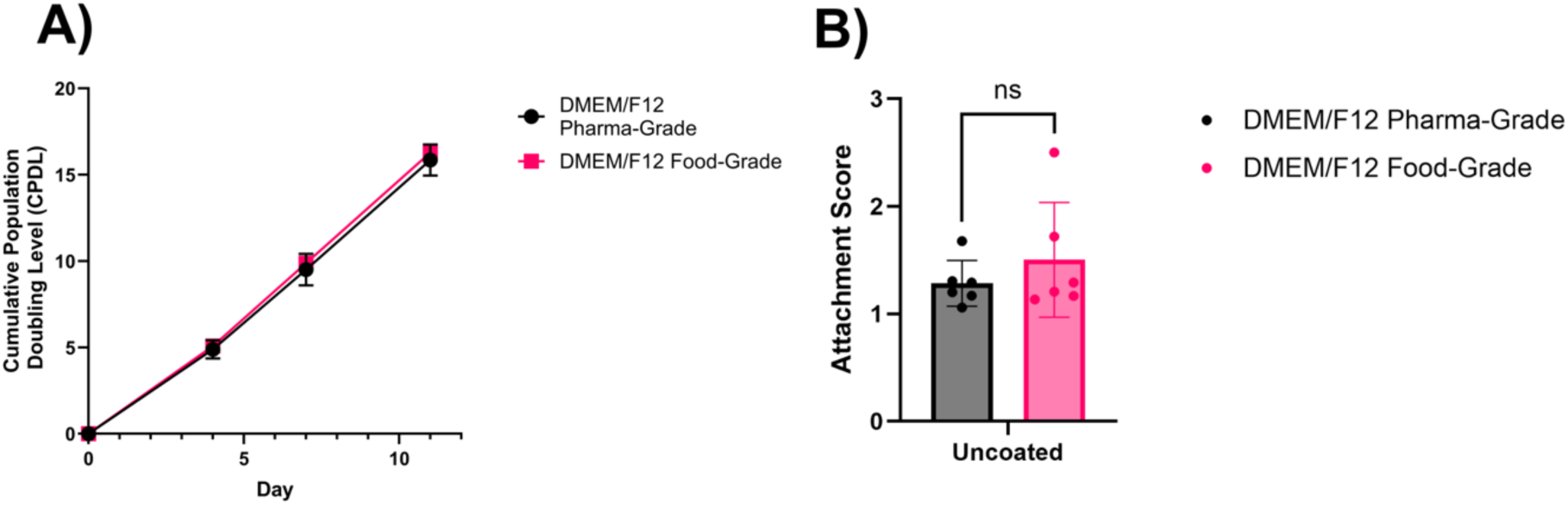
Comparison of C2C12 murine myoblast growth in food-grade vs pharmaceutical-grade DMEM/F12 basal medium when cultured in the MMM1 serum-free formulation. (A) Cumulative Population Doubling Level (CPDL) over 10 days of culture; no significant difference in growth rate was observed. Data shown as mean ± SEM (n=3). (B) Attachment efficiency on uncoated culture surfaces. Attachment scores were comparable between conditions. Data shown as mean ± SD (n=3). Significance was determined using a t-test (*p < 0.05, ns = not significant).

To substantiate the economic viability of the MMM1 formulation, a cost analysis was conducted to estimate the laboratory-scale raw material costs per litre of media (**Table 3**). Comparing modern serum-free media is often complicated by the high cost of pharmaceutical-grade recombinant human serum albumin (hSA); as the most abundant protein in serum, significant costs are associated with replacing it directly. For instance, formulations such as Beefy-9 (Stout, Mirliani, et al., 2022) and (Kolkmann et al., 2022) scale to approximately £161 and over £3,114 per litre respectively at lab-scale. The MMM1 formulation (incorporating 0.4 mg/mL hSA) similarly scaled to over £124 per litre, exceeding by a small margin the cost of standard medium containing 10% FBS (£123.79/L). Options such as plant-derived alternatives to recombinant albumin (Stout, Rittenberg, et al., 2022) which have very low raw material costs of ∼£0.001/L, could represent a solution to cost reduction in formulations where albumin is the primary cost driver.

**Table 3:**
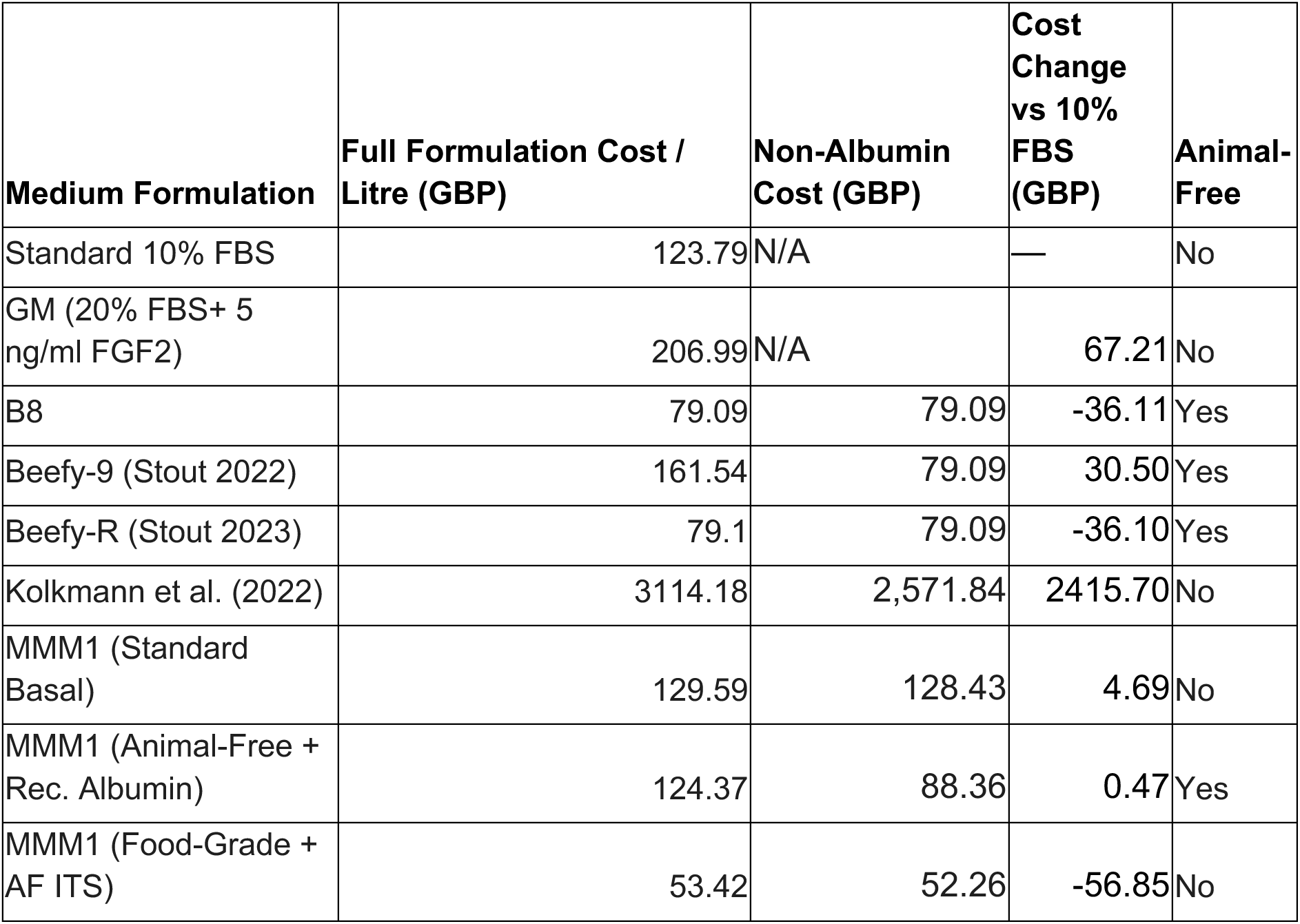
Cost analysis of lab-scale costs for highly-cited satellite cell medium formulations. Costs are estimated per litre in GBP (£) based on laboratory-scale retail pricing of individual components. The Full Formulation Cost represents the complete medium as published or optimised, including all specific growth factors, carrier proteins, and extracts. To provide a biologically and economically equitable comparison, the Non-Albumin Cost isolates the core formulation by excluding the cost of albumin a major cost driver that artificially inflates formulations costs. The Cost Change vs 10% FBS represents the percentage difference in the full formulation cost relative to the standard 10% FBS baseline. The Animal-Free column denotes the strict absence of animal-derived components.

Even when excluding the cost of albumin to ensure a fair comparison, the final MMM1 formulation proved highly cost-competitive. By using bulk food-grade DMEM/F12 (£7.50/L) alongside low-cost spirulina extract (£0.25/L), methylcellulose, and an optimised animal-free ITS blend, the final MMM1 medium achieved a cost of £53.42/L. This represents a 56.8% cost reduction compared to standard 10% FBS media.

Furthermore, these costs could be further reduced through supply chain optimisations and economies of scale. For instance, in the MMM1 formulation, beta-carotene and recombinant Hepatocyte Growth Factor (HGF) account for nearly £12.00/L of the final cost purely due to small-batch purchasing. Procuring bulk recombinant HGF (e.g., 1 mg quantities) would reduce its unit cost by over 75% compared to standard 10 µg laboratory vials. Furthermore, transitioning from highly purified laboratory-grade beta-carotene to readily available food-grade bulk equivalents (costing as little as £53.20 per kilogram) effectively eliminates its contribution to the volumetric cost. At commercial production volumes, the absolute cost of the entire formulation will drop by orders of magnitude due to the economies of scale. Bulk procurement of powdered basal media, rather than pre-hydrated liquid volumes, alongside business-to-business wholesale agreements for recombinant growth factors, effectively circumvents the massive markups associated with laboratory distribution. Consequently, while the ∼£53/L achieved by the food-grade MMM1 formulation demonstrates a significant relative cost reduction at the bench scale, it represents a conservative upper bound. Integrating this formulation into pilot-scale supply chains, where food-grade inputs like spirulina extract, methylcellulose, and beta-carotene are sourced by the metric ton, is expected to drive the final volumetric cost exponentially closer to the economic threshold required for commercial parity with conventional meat.

## 4. Limitations of the Study

While this study establishes an effective proof-of-concept framework for a cost-reduced, serum-free medium, several limitations must be acknowledged to guide future translational research.

While C2C12 serves as a reproducible model cell line for upstream screening and microcarrier bioprocess optimisation, key metabolic and physiological differences exist between spontaneously immortalised murine lines and primary livestock satellite cells. However, a trend towards the preference of immortalised cell lines rather than primary livestock cells is emerging in cultivated meat manufacturing in the products that have received regulatory approval for both human and pet consumption suggesting that the C2C12 cell line may have utility in its modelling of the cell growth characteristics of immortalised cells (Good Meat et al., 2022; Schulze, 2021). Target livestock species may display different nutritional requirements or higher sensitivity to serum withdrawal (Cosenza et al., 2022; Skrivergaard et al., 2023). Consequently, the MMM1 formulation should be viewed as a baseline proof-of-concept platform requiring species-specific optimisation.

While MMM1 successfully incorporates food-grade inputs (spirulina and methylcellulose) and a food-grade basal medium, it is important to note that the formulation is not yet entirely food-grade. The requisite growth factors, specifically FGF2 and TGF-β, currently remain high-cost, pharmaceutical-grade recombinant proteins produced via traditional *E. coli* or mammalian expression systems (Poudel et al., 2019; Savyon et al., 2025; Zhang et al., 2010). To achieve a 100% food-grade formulation, future research must focus on transitioning the production of these specific signalling proteins to high-yield, lower-cost platforms such as plant molecular farming and alternative microbial expression systems. Alternatively, cell lines may be engineered to autologously produce their own growth factors (Stout et al., 2024). Furthermore, standardising downstream processing (DSP) to food-grade filtration and purification standards, rather than highly rigorous pharmaceutical-grade chromatography, will be critical to resolving this final bottleneck (Buyel, 2025; Post, 2025).

Moreover, while the integration of bulk food-grade raw materials (such as DMEM/F12 and spirulina extract) offers significant cost reductions, it also introduces new scale-up and supply chain challenges. As demonstrated by our batch testing (**Supplementary Figure S2**), food-grade raw materials are subject to inherent batch-to-batch variability and may contain trace impurities, such as heavy metals or endotoxins, which could impact long-term cell health. To mitigate this variability at an industrial scale, biomanufacturers will need to adopt stringent lot-testing protocols. Much like the traditional lot-validation processes used for Foetal Bovine Serum (FBS), commercial facilities must identify highly effective agricultural batches and procure them in massive bulk quantities to maintain consistency across prolonged manufacturing runs. Additionally, while the spirulina extract effectively supported cell growth, its precise mechanism of action and its complex synergistic interactions with other medium components remain incompletely characterised.

## 5. Conclusions and Future Perspectives

This study successfully developed MMM1, an effective serum-free medium for cultivating C2C12 murine myoblasts. By initially screening the established B8 and Beefy-9 formulations, the study identified FGF2 and TGF-β as the most important baseline components driving cell proliferation. The novel formulation then strategically incorporated additional food-grade ingredients and growth factors to further enhance cell growth. Hepatocyte Growth Factor (HGF) significantly promoted early-stage myoblast division. Spirulina extract acted as a complex nutrient source. Additionally, food-grade methylcellulose positively influenced proliferation. Consequently, the optimised serum-free medium formulation named here as “MMM1” achieved cumulative population doublings comparable to serum-supplemented media across multiple passages without requiring cellular adaptation. MMM1 also demonstrated excellent translational potential by supporting cell expansion on gelatin-based microcarriers in suspension culture in spinner flasks. Finally, integrating a food-grade basal medium maintained cellular attachment and growth kinetics while offering a strategy to substantially reduce media input costs.

While the C2C12 cell line serves as a robust research model for bioprocessing for cultivated meat or for pet food applications, future work must translate these findings to myosatellite cells from other meat relevant species such as bovine, porcine, or avian lines. This may require adaptation to species-specific requirements; however it acts as a low cost-food grade base for targeted optimisation. As the cultivated meat industry approaches commercialisation, testing the MMM1 formulation in larger pilot-scale bioreactors will be necessary to confirm its industrial scalability. Furthermore, evaluating how these food-grade media inputs impact regulatory compliance and the nutritional profile of the final cultivated tissue will be essential for long-term viability.

## Acknowledgements

We would like to thank Leo Carillo of UCL Biochemical Engineering for his support in RT-qPCR analysis. This work was supported by the UCL EPSRC Doctoral Training Program [EP/R513143/1] and the EPSRC Cellular Agriculture Manufacturing Hub (CARMA) [EP/X038114/1].

