## Supplementary Materials for "Serum-free media development and validation for cultivation of C2C12 immortalised murine myosatellite cell line for cultivated meat"

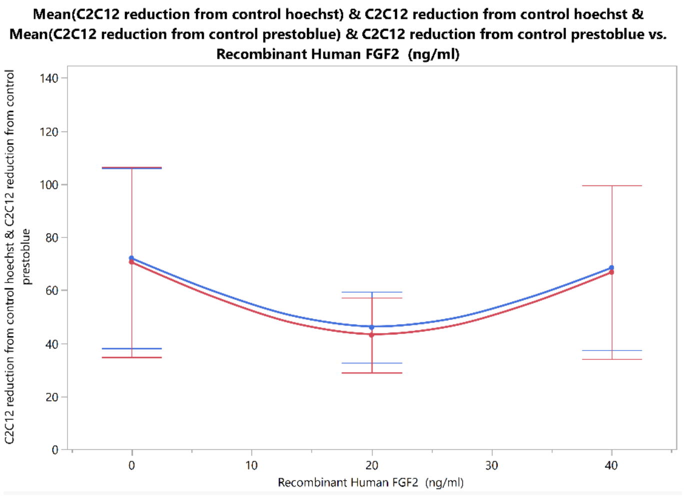


*Supplementary Figure S1:* Curvature of FGF2 Presto Blue and Hoechst

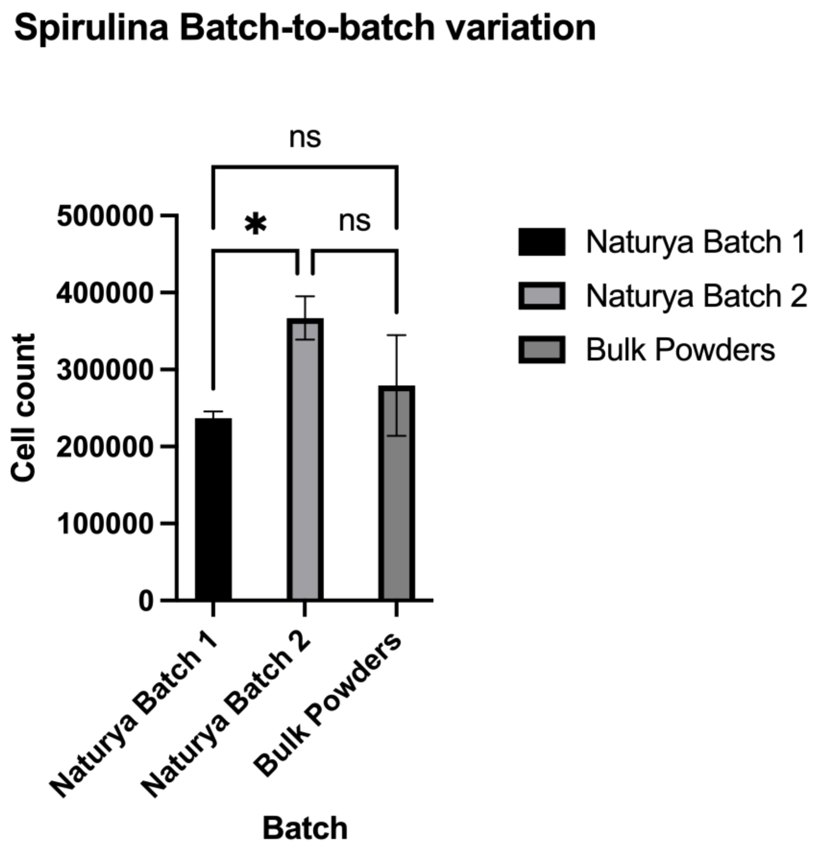


*Supplementary Figure S2: Assessment of spirulina extract batch-to-batch variability. C2C12 cell counts following culture with three distinct commercial batches of spirulina extract (Naturya 2026, Naturya 2024, and Bulk Powders). A significant difference in*

*efficacy was observed between the two Naturya batches ($p < 0.05$), highlighting the inherent variability of food-grade agricultural inputs. Data shown as mean ± SD. Statistical significance was determined using a one-way ANOVA with Tukey's multiple comparisons test (ns = not significant).*

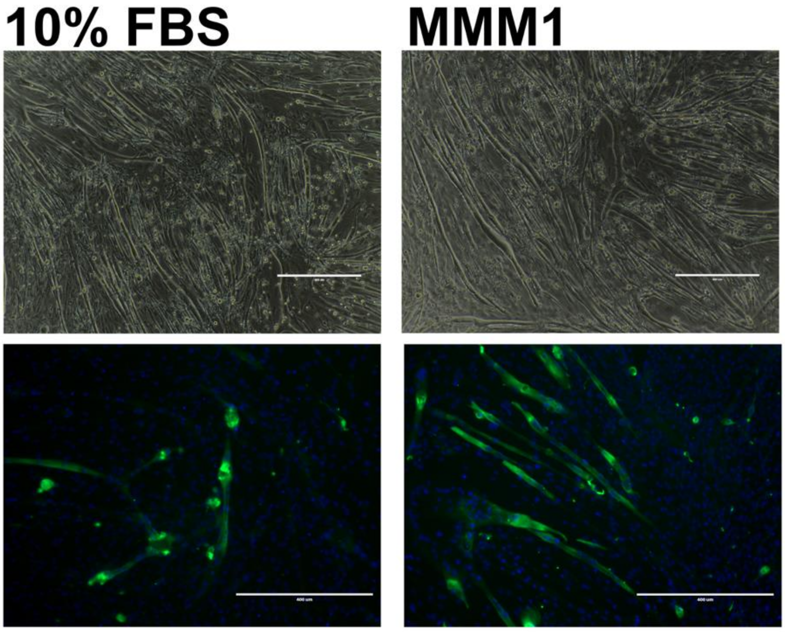


*Supplementary Figure S3: Differentiation images of C2C12s cultured in 10% FBS or MMM1 and then seeded at 30,000 cells/cm2 and differentiated for 9 days. Top row, phase contrast images, bottom row ICC images nuclei (blue) and Desmin (Green).*

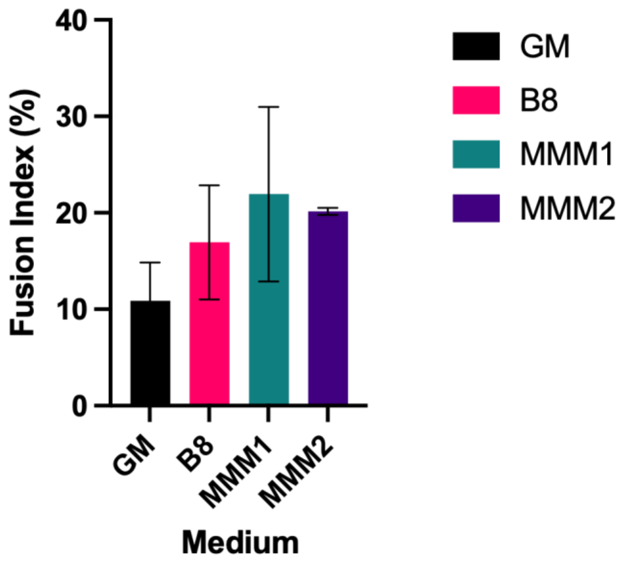


*Supplementary Figure S4: Fusion index as measured by percentage of myotubes (desmin) with a minimum of 3 nuclei per myotube (n=3). C2C12s were cultured in 10% FBS, or MMM1 and then seeded at 30,000 cells/cm2 and differentiated for 9 days. Data Shown as mean* ± SD. No significant differences observed.

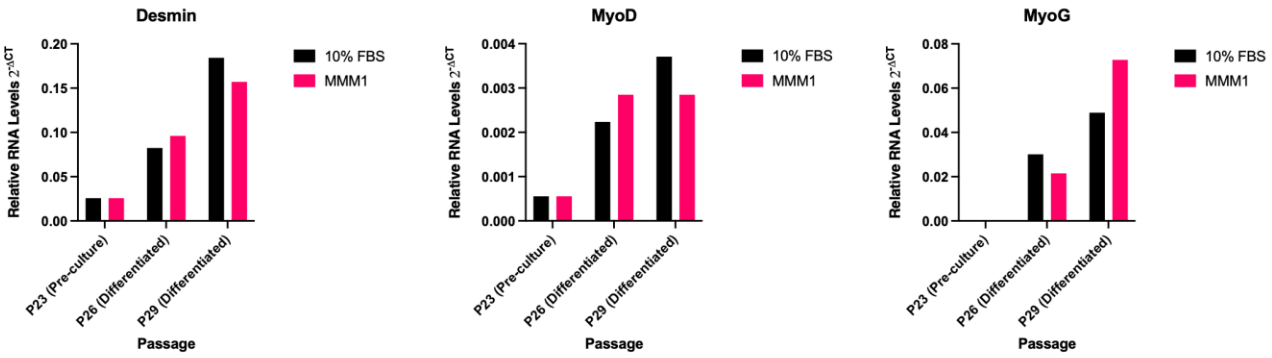


*Supplementary Figure S5: Relative expression of myogenic markers during bovine muscle stem cell culture and differentiation. Gene expression analysis of the structural protein Desmin (DES) and myogenic regulatory factors MyoD (MYOD1) and MyoG (MYOG) was performed using quantitative real-time PCR (RT-qPCR). Bovine muscle stem cells were evaluated at a pre-culture baseline (Passage 23) and following differentiation protocols at later passages (Passage 26 and Passage 29). Expression levels are presented as $2^{-\Delta CT}$ values, which have been normalised to the expression of the housekeeping gene GAPDH. Bars represent the mean of technical replicates (N=2)*

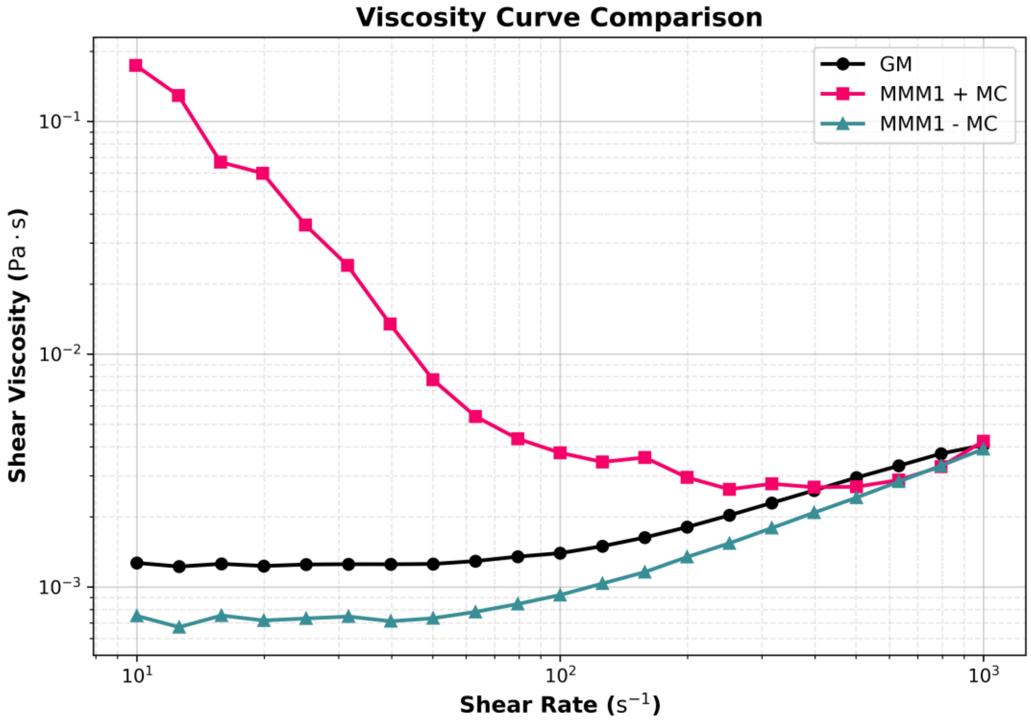


*Supplementary Figure S6: Steady-state rheological profiling of media formulations. Shear viscosity ($\eta$, Pa·s) measured as a function of shear rate ($\dot{\gamma}$, $s^{-1}$). The unsupplemented baseline (MMM1 - MC, blue triangles) and standard serum-supplemented growth medium (GM, black circles) exhibit relatively flat, near-water viscosity profiles at low-to-moderate shear rates. In contrast, the addition of methylcellulose (MMM1 + MC, pink squares) imparts a pronounced non-Newtonian, shear-thinning behaviour, characterised by significantly elevated viscosity at low shear rates that rapidly decreases and converges with the baseline controls under high shear stress ($10^3$ $s^{-1}$). This shear-responsive profile indicates that methylcellulose provides dynamic hydrodynamic shear protection in agitated culture environments while facilitating localised macromolecular crowding effects in static or protected micro-environments.*

*Supplementary Table 1: Concentration ranges used in the first definitive screening design of experiments.*

| Condition | *L-ascorbic acid 2-phosphate (mg/ml)* | *Insulin, Human Recombinant (ng/ml)* | *Transferrin human, Recombinant (ng/ml)* | *Recombinant Human FGF2 (ng/ml)* | *Sodium selenite (ng/ml)* | *NRG1, Recombinant Human (ng/ml)* | *TGFB1, Recombinant Human (ng/ml)* | *Albumin recombinant human (mg/ml)* |
|---|---|---|---|---|---|---|---|---|
| 1 | 0 | 0 | 0 | 40 | 0 | 0.1 | 0 | 0.8 |
| 2 | 0.2 | 5000 | 0 | 40 | 20 | 0 | 0 | 0.4 |
| 3 | 0 | 5000 | 2500 | 40 | 0 | 0 | 0.1 | 0.8 |
| 4 | 0.2 | 0 | 5000 | 40 | 0 | 0.05 | 0.1 | 0 |
| 5 | 0.2 | 0 | 5000 | 40 | 10 | 0 | 0 | 0.8 |
| 6 | 0.2 | 2500 | 0 | 0 | 0 | 0 | 0.1 | 0.8 |
| 7 | 0.2 | 5000 | 0 | 40 | 0 | 0.1 | 0.05 | 0 |
| 8 | 0 | 2500 | 5000 | 40 | 20 | 0.1 | 0 | 0 |
| 9 | 0.2 | 5000 | 5000 | 0 | 20 | 0 | 0.1 | 0 |
| 10 | 0.2 | 0 | 0 | 20 | 20 | 0.1 | 0.1 | 0.8 |
| 11 | 0 | 0 | 5000 | 0 | 0 | 0.1 | 0.1 | 0.4 |
| 12 | 0 | 5000 | 0 | 0 | 10 | 0.1 | 0.1 | 0 |
| 13 | 0.2 | 5000 | 5000 | 0 | 0 | 0.1 | 0 | 0.8 |
| 14 | 0 | 5000 | 5000 | 20 | 0 | 0 | 0 | 0 |
| 15 | 0 | 0 | 5000 | 0 | 20 | 0 | 0.05 | 0.8 |
| 16 | 0 | 0 | 0 | 40 | 20 | 0 | 0.1 | 0 |
| 17 | 0.1 | 5000 | 5000 | 40 | 20 | 0.1 | 0.1 | 0.8 |
| 18 | 0 | 5000 | 0 | 0 | 20 | 0.05 | 0 | 0.8 |
| 19 | 0.1 | 2500 | 2500 | 20 | 10 | 0.05 | 0.05 | 0.4 |
| 20 | 0.1 | 0 | 0 | 0 | 0 | 0 | 0 | 0 |
| 21 | 0.2 | 0 | 2500 | 0 | 20 | 0.1 | 0 | 0 |

*Supplementary Table 2: Component concentration ranges of screen 2, all cultures were supplemented with ITSX (10 μg/ml Insulin, 5.5 μg/ml Transferrin, 6.7 ng/ml Sodium selenite, 2 μg/ml Ethanolamine)*

| Condition | HGF, Recombinant, Human (ng/ml) | IGF-1, Recombinant, Human (ng/ml) | PDGF-BB, Recombinant, Human (ng/ml) | VEGF Recombinant, Human (ng/ml) | IL-6, Recombinant, Human (ng/ml) | Recombinant Human FGF2 (ng/ml) | TGFB1, Recombinant Human (ng/ml) | NRG-1, Recombinant, Human (ng/ml) | Albumin, Recombinant, Human (mg/ml) |
|---|---|---|---|---|---|---|---|---|---|
| 1 | 0 | 5 | 1 | 0 | 1 | 20 | 0 | 0.2 | 0 |
| 2 | 0 | 5 | 0.5 | 0 | 1 | 0 | 0 | 0 | 40000 |
| 3 | 0 | 2.5 | 0 | 1 | 0 | 0 | 0 | 0.2 | 40000 |
| 4 | 0 | 5 | 0 | 1 | 0.5 | 0 | 0.2 | 0 | 0 |
| 5 | 0.25 | 5 | 1 | 1 | 0 | 0 | 0 | 0.2 | 0 |
| 6 | 0.25 | 0 | 1 | 1 | 1 | 0 | 0 | 0 | 40000 |
| 7 | 0.25 | 0 | 0.5 | 1 | 0 | 40 | 0.2 | 0.2 | 0 |
| 8 | 0.125 | 2.5 | 0.5 | 0.5 | 0.5 | 20 | 0.1 | 0.1 | 20000 |
| 9 | 0 | 5 | 0 | 0 | 0 | 40 | 0.2 | 0.2 | 0 |
| 10 | 0.25 | 0 | 1 | 0 | 0.5 | 40 | 0 | 0.2 | 40000 |
| 11 | 0.25 | 5 | 0 | 0 | 0 | 40 | 0 | 0.1 | 40000 |
| 12 | 0 | 0 | 0 | 0 | 1 | 40 | 0.2 | 0 | 40000 |
| 13 | 0.25 | 0 | 0 | 0 | 1 | 0 | 0.1 | 0.2 | 0 |
| 14 | 0.25 | 5 | 1 | 0 | 0 | 0 | 0.2 | 0 | 20000 |
| 15 | 0 | 0 | 1 | 1 | 1 | 0 | 0.2 | 0.1 | 0 |
| 16 | 0 | 0 | 0 | 1 | 1 | 40 | 0 | 0.2 | 20000 |
| 17 | 0.25 | 5 | 0 | 0.5 | 1 | 0 | 0.2 | 0.2 | 40000 |
| 18 | 0 | 0 | 1 | 0.5 | 0 | 40 | 0 | 0 | 0 |
| 19 | 0 | 5 | 1 | 1 | 0 | 40 | 0.1 | 0 | 40000 |
| 20 | 0 | 0 | 1 | 0 | 0 | 0 | 0.2 | 0.2 | 40000 |
| 21 | 0.125 | 5 | 1 | 1 | 1 | 40 | 0.2 | 0.2 | 40000 |
| 22 | 0.25 | 0 | 0 | 1 | 0 | 20 | 0.2 | 0 | 40000 |
| 23 | 0.25 | 5 | 0 | 1 | 1 | 40 | 0 | 0 | 0 |
| 24 | 0.125 | 0 | 0 | 0 | 0 | 0 | 0 | 0 | 0 |
| 25 | 0.25 | 2.5 | 1 | 0 | 1 | 40 | 0.2 | 0 | 0 |

*Supplementary Table 3: Definitive screening design, with base medium ITSXF. : Alb: Albumin, Recombinant, Human; HGF: Hepatocyte Growth Factor; TGF: Transforming*

*Growth Factor; PDGF: Platelet-Derived Growth Factor-BB; IGF1: Insulin-like Growth Factor-1; IL6: Interleukin-6; B-Car: Beta Carotene; Curc: Curcumin; Ala: Raecemic alanine; Carn: L-carnosine; MC: Methylcellulose; Spir: Spirulina extract; HC: Hydrocortisone; LA: Linoleic acid.*

| **Condition** | **Alb (mg/ml)** | **HGF (ng/ml)** | **TGF (ng/ml)** | **PDGF (ng/ml)** | **IGF-1 (ng/ml)** | **IL-6 (ng/ml)** | **B-Car (mg/ml)** | **Curc (mg/mL)** | **Ala (mg/mL)** | **Carn (mg/mL)** | **MC (mg/mL)** | **Spir % v/v** | **HC (μg/mL)** | **LA (μg/mL)** |
|---|---|---|---|---|---|---|---|---|---|---|---|---|---|---|
| 1 | 0 | 0 | 0 | 0 | 0 | 2 | 0 | 0.01 | 0.02 | 0 | 0.1 | 0 | 0 | 1 |
| 2 | 0.4 | 0 | 0.4 | 2 | 0 | 2 | 0.001 | 0.01 | 0 | 0.2 | 0 | 0.2 | 0 | 0 |
| 3 | 0.4 | 0 | 0 | 0 | 10 | 0 | 0 | 0.01 | 0 | 0.2 | 0.1 | 0.2 | 0.2 | 0.1 |
| 4 | 0 | 0 | 0.4 | 0 | 10 | 0 | 0.001 | 0 | 0 | 0.2 | 0 | 0 | 0.2 | 1 |
| 5 | 0.4 | 0.5 | 0 | 0 | 10 | 0 | 0.001 | 0 | 0.02 | 0 | 0 | 0 | 0 | 0 |
| 6 | 0 | 0 | 0 | 0 | 10 | 2 | 0 | 0.01 | 0 | 0 | 0 | 0 | 0.2 | 0 |
| 7 | 0 | 0.5 | 0 | 2 | 10 | 2 | 0.001 | 0.01 | 0.02 | 0.2 | 0 | 0 | 0.02 | 1 |
| 8 | 0.4 | 0.5 | 0.4 | 2 | 10 | 2 | 0.001 | 0 | 0 | 0 | 0.1 | 0.2 | 0.2 | 1 |
| 9 | 0 | 0 | 0.4 | 0 | 10 | 0 | 0.001 | 0 | 0.02 | 0 | 0 | 0.2 | 0.2 | 0.1 |
| 10 | 0.4 | 0.5 | 0 | 0 | 0 | 2 | 0.001 | 0 | 0.02 | 0.2 | 0.1 | 0 | 0.2 | 0 |
| 11 | 0.4 | 0 | 0.4 | 2 | 0 | 2 | 0 | 0 | 0.02 | 0 | 0 | 0 | 0.02 | 0.1 |
| 12 | 0 | 0 | 0.4 | 2 | 0 | 2 | 0 | 0 | 0.02 | 0 | 0.1 | 0.2 | 0.2 | 0 |
| 13 | 0 | 0 | 0.4 | 0 | 10 | 2 | 0.001 | 0 | 0.02 | 0.2 | 0.1 | 0.2 | 0 | 1 |
| 14 | 0 | 0.5 | 0.4 | 2 | 10 | 0 | 0 | 0.01 | 0 | 0 | 0.1 | 0 | 0 | 0 |
| 15 | 0 | 0.5 | 0 | 0 | 0 | 2 | 0.001 | 0.01 | 0 | 0 | 0 | 0.2 | 0.2 | 0.1 |
| 16 | 0 | 0.5 | 0.4 | 0 | 10 | 2 | 0 | 0 | 0 | 0.2 | 0.1 | 0 | 0.02 | 0.1 |
| 17 | 0.4 | 0 | 0.4 | 0 | 0 | 0 | 0 | 0.01 | 0.02 | 0.2 | 0.1 | 0 | 0.2 | 1 |
| *18* | *0.4* | *0.5* | *0.4* | *2* | *10* | *0* | *0.001* | *0.01* | *0.02* | *0* | *0.1* | *0* | *0.2* | *0.1* |
| *19* | *0* | *0* | *0* | *2* | *0* | *0* | *0.001* | *0* | *0* | *0.2* | *0.1* | *0* | *0* | *0.1* |
| *20* | *0.4* | *0.5* | *0* | *0* | *0* | *0* | *0* | *0* | *0* | *0* | *0.1* | *0.2* | *0* | *1* |
| *21* | *0.4* | *0.5* | *0.4* | *0* | *10* | *2* | *0* | *0.01* | *0.02* | *0.2* | *0* | *0.2* | *0* | *0.1* |

| | | | | | | | | | | | | | | |
|---|---|---|---|---|---|---|---|---|---|---|---|---|---|---|
| *22* | *0.4* | *0* | *0.4* | *0* | *0* | *0* | *0.001* | *0.01* | *0* | *0* | *0* | *0* | *0.02* | *1* |
| *23* | *0.4* | *0.5* | *0.4* | *2* | *0* | *2* | *0* | *0* | *0* | *0.2* | *0* | *0* | *0.2* | *1* |
| *24* | *0.4* | *0* | *0* | *2* | *10* | *2* | *0.001* | *0.01* | *0* | *0* | *0.1* | *0.2* | *0.02* | *1* |
| *25* | *0* | *0.5* | *0* | *2* | *0* | *0* | *0* | *0.01* | *0.02* | *0.2* | *0* | *0.2* | *0.2* | *1* |
| *26* | *0.4* | *0* | *0* | *2* | *10* | *0* | *0* | *0* | *0.02* | *0.2* | *0* | *0.2* | *0.02* | *0* |

*Supplementary Table 4: Comparison of contents of B8 (Kuo et al., 2020), Beefy-9 (Stout et al., 2022), Kolkmann (Kolkmann et al., 2022), (Skrivergaard et al., 2023), and MMM1 components*

| **Component Category** | **Supplement** | **B8** | **Beefy-9** | **Kolkmann et al. (2022)** | **Skrivergaard et al. (2023)** | **MMM1 (This Study)** |
|---|---|---|---|---|---|---|
| **Basal Medium** | DMEM/F12 | ✓ | ✓ | ✓ | ✓ | ✓ *(Food-Grade optional)* |
| | GlutaMAX™ | — | — | ✓ | — | ✓ |
| **Carrier Proteins & Extracts** | Albumin | — | ✓ *(Rec. HSA)* | ✓ *(Rec. HSA)* | ✓ *(BSA)* | ✓ *(BSA or Rec. HSA)* |
| | Spirulina Extract | — | — | — | — | ✓ |
| | Fibronectin | — | — | ✓ | — | — |
| | Fetuin | — | — | — | ✓ | — |
| **Core Metabolism (ITS)** | Insulin | ✓ | ✓ | ✓ *(via ITS-X)* | ✓ *(via ITS)* | ✓ *(via ITS-X or AF-ITS)* |
| | Transferrin | ✓ | ✓ | ✓ *(via ITS-X)* | ✓ *(via ITS)* | ✓ *(via ITS-X or AF-ITS)* |
| | Sodium Selenite | ✓ | ✓ | ✓ *(via ITS-X)* | ✓ *(via ITS)* | ✓ *(via ITS-X or AF-ITS)* |
| **Growth Factors** | FGF-2 | ✓ | ✓ | ✓ | ✓ | ✓ |
| | TGF-beta | ✓ | ✓ | — | — | ✓ |
| | NRG-1 | ✓ | ✓ | — | — | — |
| | HGF | — | — | ✓ | — *(✓ optional)* | ✓ |
| | PDGF-BB | — | — | — | — *(✓ optional)* | ✓ |
| | IGF-1 | — | — | ✓ | — | — |
| | VEGF | — | — | ✓ | — | — |

| | | | | | | |
|---|---|---|---|---|---|---|
| | IL-6 | — | — | ✓ | — | — |
| **Small Molecules & Polymers** | L-Ascorbic Acid 2-phosphate | ✓ | ✓ | ✓ | — | — |
| | Methylcellulose | — | — | — | — | ✓ |
| | Beta-Carotene | — | — | — | — | ✓ |
| | Hydrocortisone | — | — | ✓ | — | — |
| | alpha-linolenic acid | — | — | ✓ | — | — |

*Supplementary Table 5: DoE Definitive screening design for optimisation of animal-free and high use growth factors.*

| | **Albumin** | **ITS** | **FGF2 (ng/ml)** | **TGF-B (ng/ml)** |
|---|---|---|---|---|
| 1 | Recomb | Optimal | 40 | 0.4 |
| 2 | None | AF | 20 | 0.4 |
| 3 | Recomb | AF | 20 | 0.1 |
| 4 | Recomb | AF | 30 | 0 |
| 5 | BSA | Animal | 40 | 0.1 |
| 6 | None | Optimal | 30 | 0.1 |
| 7 | None | Animal | 40 | 0 |
| 8 | Recomb | Animal | 30 | 0 |
| 9 | None | AF | 40 | 0 |
| 10 | None | Optimal | 30 | 0.1 |
| 11 | BSA | Optimal | 20 | 0 |
| 12 | Recomb | Optimal | 40 | 0.4 |
| 13 | Non | Optimal | 20 | 0 |
| 14 | Non | Animal | 30 | 0.4 |
| 15 | Recomb | Animal | 20 | 0.1 |
| 16 | BSA | AF | 30 | 0.4 |
| 17 | BSA | AF | 40 | 0.1 |
| 18 | None | Animal | 20 | 0.4 |
